# Training and spontaneous reinforcement of neuronal assemblies by spike timing plasticity

**DOI:** 10.1101/066969

**Authors:** Gabriel Koch Ocker, Brent Doiron

## Abstract

The synaptic connectivity of cortex is plastic, with experience shaping the ongoing interactions between neurons. Theoretical studies of spike timing–dependent plasticity (STDP) have focused on either just pairs of neurons or large-scale simulations. A simple analytic account for how fast spike time correlations affect both micro- and macroscopic network structure is lacking. We develop a low-dimensional mean field theory for STDP in recurrent networks and show the emergence of assemblies of strongly reciprocally coupled neurons with shared stimulus preferences. After training this connectivity is actively reinforced by spike train correlations during the spontaneous dynamics. Furthermore, the stimulus coding by cell assemblies is actively maintained by these internally generated spiking correlations, suggesting a new role for noise correlations in neural coding. Assembly formation has been often associated with firing rate-based plasticity schemes; our theory provides an alternative and complementary framework, where fine temporal correlations and STDP form and actively maintain learned structure in cortical networks.

## Introduction

A cornerstone principle that bridges systems and cellular neuroscience is that the synaptic wiring between neurons is sculpted by experience. The early origins of this idea are often attributed to Donald Hebb [Hebb, 1949, Markram et al., 2011, Harris and Mrsic-Flogel, 2013], who famously postulated that groups of neurons that are repeatedly coactivated will strengthen their synaptic wiring. The interconnected group, termed an *assembly*, has become an essential building block of many theories of neural computation [Buzski, 2010] and associative memory [Neves et al., 2008].

Despite the functional appeal of neuronal assemblies, physiological evidence of assembly structure has only recently been collected. In rodent sensory cortices, pyramidal neurons have stimulus-specific connectivity and connectivity strength [Ko et al., 2011, Cossell et al., 2015, Lee et al., 2016], arranged in clustered architectures [Yoshimura et al., 2005, Perin et al., 2011, Lee et al., 2016]. This functional connectivity is enhanced during development and by sensory experience, suggesting activity-dependent long-term plasticity as a key mechanism for assembly formation [Ko et al., 2013, Ko et al., 2014]. Temporal correlations between pre- and postsynaptic activity can be a crucial determinant of long-term plasticity [Markram et al., 2012]. Hebbian spike timing–dependent plasticity (STDP) reinforces temporally causal interactions, while weakening synapses not involved in causing postsynaptic spikes. Modeling and theoretical studies have shown than Hebbian STDP promotes the development of feedforward architectures [Masuda and Kori, 2007, Takahashi et al., 2009, Tannenbaum and Burak, 2016] giving rise to sequential activity [Gerstner et al., 1993, Fiete et al., 2010]. In contrast, the role of Hebbian STDP in assembly formation remains unclear.

The amplitude of potentiation and depression in STDP depends on pre- and postsynaptic firing rates [Sjstrm et al., 2001]. Models with rate-dependent STDP admit reductions to classic rate-based plasticity rules [Pfister and Gerstner, 2006, Clopath, Claudia et al., 2010] so that when high (low) postsynaptic activity is paired with high presynaptic activity, synaptic connections are potentiated (depressed) (Fig. 1A). This rate-based plasticity can give rise to stable assembly structure [Mongillo et al., 2005, Litwin-Kumar and Doiron, 2014, Zenke et al., 2015] via firing rate transitions that toggle between strongly potentiation- and depression-dominated regimes (Fig. 1B). In such training protocols, fast spike time correlations contribute minimally to plasticity.

**Figure 1.**
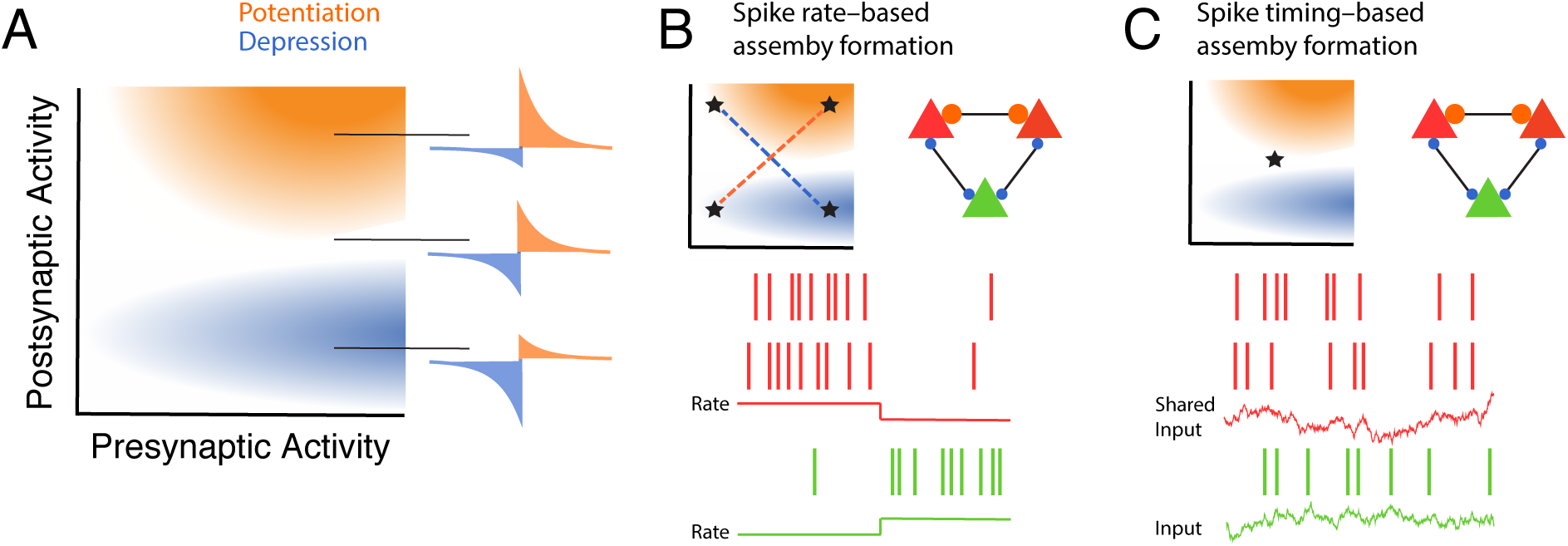
Spike rate– versus spike timing–based neuronal assembly formation. (A). Schematic illustrating how combinations of pre- and postsynaptic activity combine to drive synaptic potentiation and depression in models of STDP. The schematic is adapted from Litwin-Kumar & Doiron [Litwin-Kumar and Doiron, 2014] where STDP rules based on third-order spike interactions [Pfister and Gerstner, 2006], or voltage- [Clopath, Claudia et al., 2010] or calcium-based [Graupner and Brunel, 2012] learning were studied. The STDP curves on the right indicate the degree of potentiation and depression as pre- and postsynaptic activity ranges. (B) Example three neuron group with two neurons having co-fluctuating firing rates (bottom, red–red) and the other neuron having anti-correlated firing rate fluctuations (bottom, green–red). This dynamic potentiates synaptic coupling between correlated neurons while depressing synaptic coupling between anti-correlated neurons (right graph). (C) Same as B except firing rates are fixed at a value that balances rate-based potentiation and depression. Shared input correlations to two neurons can potentiate strong recurrent synapses (bottom, red-red) and depress uncorrelated neurons (bottom, green-red).

Spike trains in diverse cortical areas exhibit covariable trial-by-trial fluctuations (noise correlations). These noise correlations covary with neurons’ stimulus preferences and synaptic connectivity, suggesting that assembly structure and noise correlations are related [Bair et al., 2001, Kohn and Smith, 2005, Rothschild et al., 2010, Ko et al., 2011, Cossell et al., 2015]. Furthermore, excitatory-inhibitory interactions within cortical circuits create nearly synchronous temporal structure between spike trains that overlaps with the fine timescale required for STDP learning [Brgers et al., 2012, Jia et al., 2013, Salkoff et al., 2015]. The precise timing of pre- and postsynaptic spikes can be a crucial determinant of plasticity in vivo [Feldman, 2012], and recently it has been shown that fine timescale optogenetic photostimulation in cortex can form assembly structure in vivo [Kim et al., 2016, Carrillo-Reid et al., 2016]. Thus, there is sufficient experimental evidence to suggest that even when neurons’ overall firing rates are static, spike timing may play an important role in assembly formation and stability.

We show that spike time correlations can, in the absence of rate-based plasticity mechanisms, form Hebbian assemblies in response to spatially correlated external inputs in model spiking networks (Fig. 1C). We develop a low-dimensional theory revealing that parallel training of assemblies through spike timing promotes strong connectivity and strong *reciprocal* connectivity within co-stimulated groups. After training, the internally generated spike time correlations in our models reinforce learned architectures during spontaneous activity. Finally, this result motivates us to speculate on a new beneficial role of internally generated noise correlations for stimulus coding: to maintain stimulus-specific assembly wiring that supports enhanced response sensitivity.

## Results

### Plasticity of partially symmetric networks during spontaneous activity

We first present the basic network properties of our network (Methods: Network Model). We modeled networks of 1500 excitatory neurons and 300 inhibitory neurons, both types following exponential integrate-and-fire dynamics [Fourcaud-Trocme et al., 2003]. In order to focus on learning due to precise spike time correlations, we began by studying a classical Hebbian spike pair–based plasticity rule for the plasticity between excitatory neurons (eSTDP) [Gerstner et al., 1996, Markram, 1997, Bi and Poo, 1998] (Fig. 2A; Methods: Plasticity Models). The plasticity rule is phenomenological, and embodies the simple observation that spike pairs induce changes in synaptic weights and the amplitude of these changes depends on the time lag between the two spikes [Markram et al., 2012]. Inhibitory synapses likewise underwent inhibitory STDP in order to maintain stable activity [Vogels et al., 2011, Luz and Shamir, 2012, Litwin-Kumar and Doiron, 2014, Zenke et al., 2015] (Fig. 2B; Methods: Plasticity Models). The plasticity was slow compared to the timescales of the membrane dynamics and spike discharge (Fig. 2C). This separation of timescales between spike time and synaptic weight dynamics permitted an averaging theory in which individual synaptic weights evolve according the STDP rule and the frequency of pairs of pre- and postsynaptic spikes at different time lags [Kempter et al., 1999]. This frequency is given by their firing rates and the cross-covariance function of the two spike trains (Fig. 2D).

**Figure 2.**
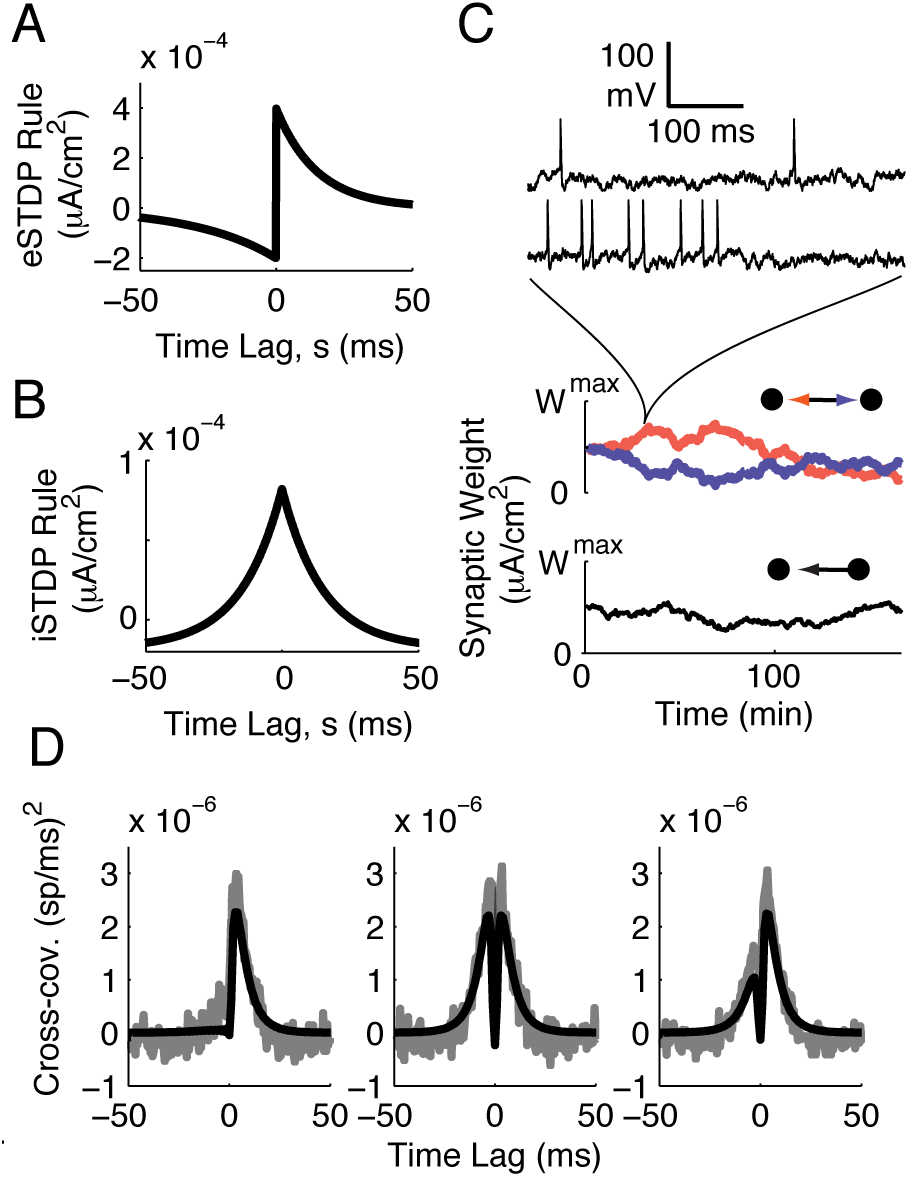
Network structure shapes synaptic plasticity. (A) The eSTDP rule, *L*(*s*), is composed of exponential windows for depression (-) and potentiation (+). Each is defined by its amplitude *f*_*±*_ and timescale *τ*_*±*_. (B) The iSTDP rule, *L*_*I*_ (*s*), is defined by its timescale *τ*_*I*_ and the amplitudes of inhibitory potentiation and postsynaptic depression, *d*_*I*_. (C) Synaptic weights evolve on a slow timescale. Individual synaptic weights are governed by the relative timing of spikes in the pre- and postsynaptic neurons’ spike trains. (D) Average spike train covariance between monosynaptically connected pairs (left), reciprocally connected pairs (center) and all pairs (right). Shaded lines: simulation. Solid lines: linear response theory (first-order truncation, Eq. S.4).

Rather than examining the plasticity of every individual synaptic weight, we took a simple characterization of the network’s excitatory-excitatory structure in terms of two variables:

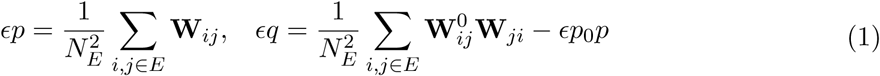

These measure the mean weight of excitatory-excitatory synapses (*p*) and the mean weight of *reciprocal* excitatory-excitatory synapses (*q*) above what would be expected in an unstructured network. In addition, we examined the plastic strength of the inhibitory-excitatory mean weight 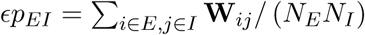. In these equations, the parameter *ϵ* = 1/(*N*_*E*_*p*_0_) scales connection strengths by the mean number of excitatory inputs to an excitatory neuron.

Starting from the plasticity of individual synaptic weights, we derived equations governing the plasticity of the averaged connectivity variables *p* and *q* (see Supporting Information S.1 for a complete derivation). In our derivation we make two main approximations. First, the average spike train covariances include contributions from only direct synaptic connections and common inputs, neglecting contributions from disynaptic and longer paths through the network. Second, we ignore the bounds on synaptic weights, meaning that our theory only describes the networks’ transient dynamics, not equilibrium states of the connectivity. These two simplifications allow us to calculate how the dynamics of the average spike train cross-correlation of connected neurons drives plasticity of the average excitatory-excitatory synaptic weight *p*, and similarly, the average spike train cross-correlation between reciprocally coupled neurons drives plasticity of the mean reciprocal synaptic weight *q*:

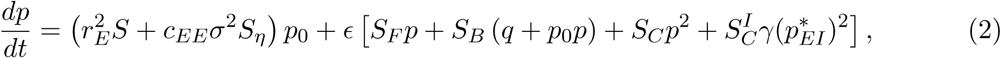

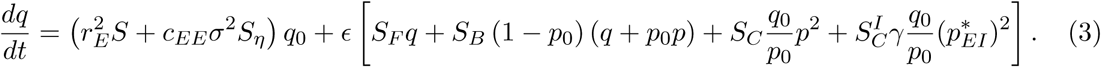

The different terms on the right-hand side of each of these equations arise from different sources of spiking correlation. The first terms on the right-hand side of (2) describe the contributions of chance spike coincidences 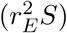, with *r*_*E*_ being the network-averaged firing rate, and of correlations induced by external inputs (*c*_*EE*_*σ*^2^*S*_*η*_). Here *S* is the total area under the eSTDP rule, determining whether STDP is potentiation-dominated or depression-dominated. Similarly, *S*_*η*_ measures the area under the STDP rule weighted by the average susceptibility of two neurons to the external input correlations (Supporting Information S.1.). The latter terms describe the contribution of correlations induced by coupling within the network, also weighted by the eSTDP rule. The effect of correlations due to direct (forward) connections is measured by *S*_*F*_, and those due to reciprocal (backward) connections is measured by *S*_*B*_. The final terms arise from correlations due to common inputs from excitatory (*S*_*C*_) or inhibitory 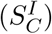 neurons. The parameter *γ* is the ratio of the number of inhibitory neurons to excitatory neurons (here *γ* = 1*/*5), and 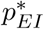 is the homeostatic fixed point of the mean inhibitory-excitatory weight, which we will elaborate on after discussing the excitatory-excitatory plasticity dynamics. The dynamics of the mean reciprocal synaptic weight, *q*_*EE*_, have a similar form to those of *p*_*EE*_. The only new parameter, *q*_0_, is the empirical frequency of reciprocal synapses in the network above chance levels. It is analogous to *q* but measured from the adjacency matrix 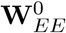 rather than the weight matrix **W**.

We assumed a balance between potentiation and depression, so that chance spike coincidences did not dominate the plasticity (star in Figure 1C). This assumption, when combined with an absence of training (*c*_*EE*_ = 0), leads to the synaptic dynamics being governed by different sources of internally generated spiking covariability, each interacting with the eSTDP rule *L*(*s*). Spiking covariations from direct connections mainly contribute at positive time lags, interacting with the potentiation side of the eSTDP rule. This is reflected in the average spike train covariance between monosynaptically connected neurons (Fig. 2D, left). Reciprocal connections, in contrast, contribute spiking covariations at negative time lags, interacting with the depression side of the eSTDP rule. This depression is seen in the average spike train covariance between reciprocally connected pairs, which includes the contributions from both direct and reciprocal connections (Fig. 2D, middle). Finally, the contributions from common inputs are temporally symmetric around zero time lag, interacting with both the potentiation and depression windows. The average spike train covariance between all neurons was asymmetric because of the presence of non-symmetric connections (Fig. 2D, right). This reduced theory provides a good quantitative prediction of the plasticity within our large-scale integrate-and-fire network (Fig. **??**B, compare the solid theory curves to the shaded curves estimated from numerical simulations). It also provides a qualitative understanding of the network dynamics.

It has been long known that additive Hebbian eSTDP produces unstable synaptic dynamics for pairs of coupled neurons through a competition between potentiation and depression [Song et al., 2000, Van Rossum et al., 2000]. Our theory extends this idea to large populations of neurons through mean field dynamics of *p* and *q*, where the different sources of spiking covariability in Eqs. (2) and (3) impose thresholds for potentiation or depression of *p* and *q*. These thresholds occur at what are mathematically called the *nullclines* of the dynamical system. A nullcline is a set of points (*p, q*) where one variable does not change; the *p*-nullcline is the collection of (*p, q*) values where *dp/dt* = 0, and similarly the *q*-nullcline is the (*p, q*) values where *dq/dt* = 0. In other words, if the network starts with *p* and *q* values that lie exactly on the *p* nullcline, then the amounts of potentiation and depression occurring amongst excitatory-excitatory synapses will be exactly balanced so that the mean synaptic weight will not change (*dp/dt* = 0); the same is true at the *q* nullcline. On either side of the nullcline, whether *dp/dt* is positive or negative dictates if *p* potentiates or depresses (and likewise for *q*). Below we will consider the plasticity dynamics the nullcline structure generates and the conditions for stable activity during plasticity.

The nullclines of Eqs. (2) and (3) intersect at a single point in (*p, q*) space, marking an equilibrium point of the system (since both *dp/dt* = 0 and *dq/dt* = 0 at the intersection). For the Hebbian plasticity rule used (Fig. 2A) the equlibrium point is an unstable repeller with dynamics flowing away from the point (Fig. **??**A, red arrows). The nullclines act as thresholds, so that if either *p* or *q* were initially stronger than its threshold it would potentiate and otherwise it would depress. The combination of the *p* and *q* nullclines then partition (*p, q*) space into four regions, where distinct evolution of *p* and *q* occur (Fig. **??**A, numbered regions). For instance, when the initial *p* and *q* values are both low (region 1 in Fig. **??**A and Fig. **??**B, left) then both the average connection strength as well as the strength of reciprocal connections depress. In contrast, when the initial *p* value is higher and lies on the other side of the *p*-nullcline (region 2 in Fig. **??**A and Fig. **??**B, right) then the average connection strength potentiates while the strength of reciprocal connections continues to depress. The low dimensional mean field dynamics of Eqs. (2) and (3) not only gives a qualitative understanding of synaptic dynamics, it also provides a good quantitative prediction of the plasticity within our large-scale integrate-and-fire network (Fig. **??**C, compare the solid theory curves to the shaded curves estimated from numerical simulations). In total, by marking the nullcline structure of Eqs. (2) and (3) we are able to predict combined dynamics of potentiation and depression of synaptic wiring in the recurrent network.

As mentioned above, we assume that potentiation and depression are balanced. To investigate the degree to which our results depend upon this balance we systematically varied the amplitudes of the depression and potentiation windows in the eSTDP rule: *F*_−_ ← *F*_−_ + *α*, *F*_+_ *← F*_+_ + *α*. For *α <* 0 this shifted the balance toward depression, and vice versa. At each *α*, we computed the area of each dynamical regions discussed above (Fig. 3D; for calculation details see Supporting Information S.3). For strong enough depression, only region 1 existed (depression for both *p* and *q*). At intermediate *α*, a region with depression for *p* but potentiation for *q* emerged. This surprising dynamics is due to the definition of *q* as the above-chance strength of reciprocal synapses. For depression-dominated plasticity, all synaptic weights depressed [Ocker et al., 2015] but reciprocal synapses did so slower than would be expected in an unstructured network. For slightly higher *α* a region with potentiation for *p* but depression for *q* emerged. As the eSTDP balance shifted more towards potentiation, these dynamical regions disappeared and only potentiation of *p* and *q* remained. Thus, while interesting *p* and *q* dynamics depend upon balance between potentiation and depression they persist as the eSTDP rule is titled away from balance to a moderate degree.

**Figure 3.**
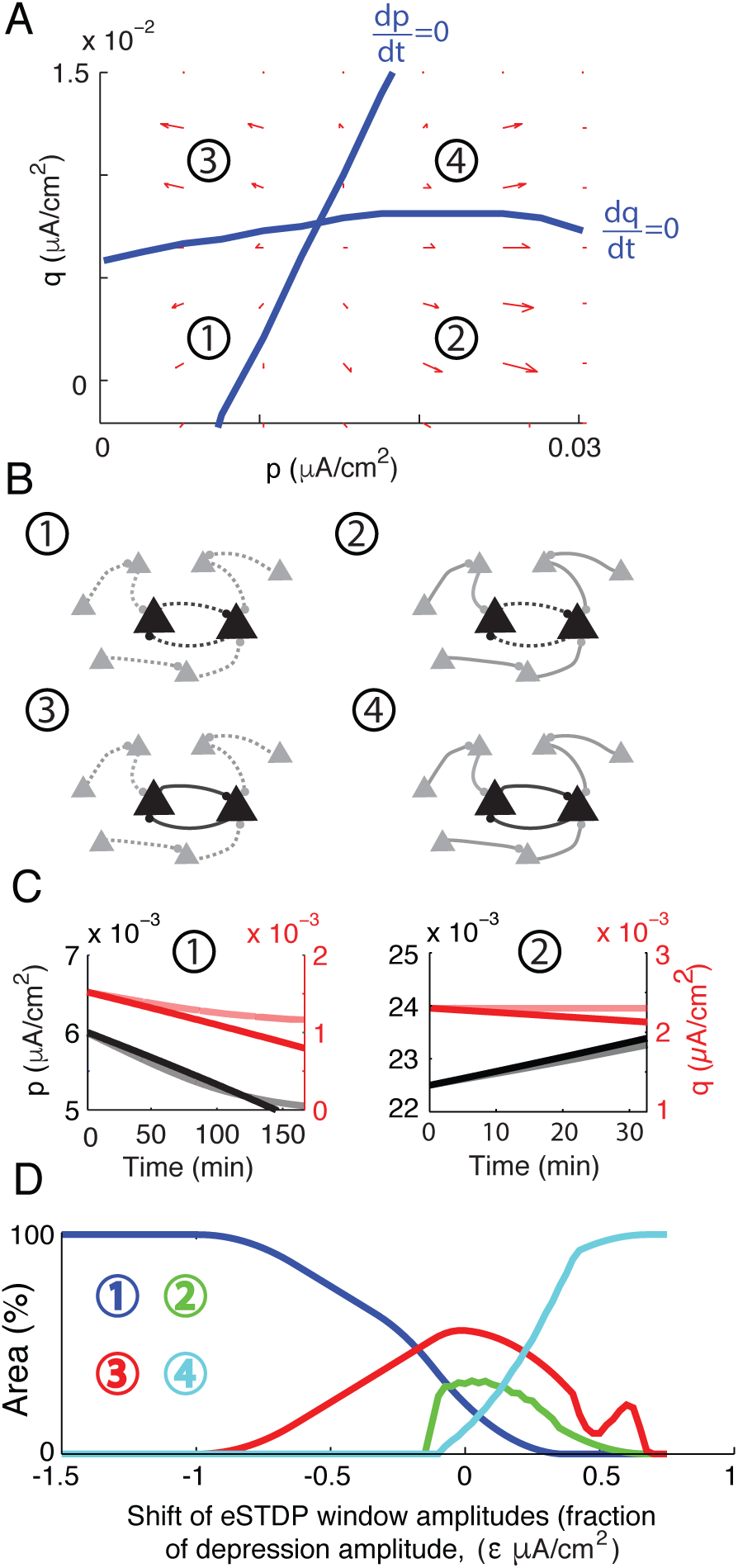
Spontaneous dynamics of the excitatory network structure. (A) Dynamics of the mean synaptic weight (*p*) and the mean above-chance strength of *reciprocal* synapses, *q*. There is a threshold for potentiation of each given by its nullcline (blue lines). These partitioned the *p, q* phase space into regions where each could potentiate or depress. (B) Illustrations of the network structures promoted in each of the four possible regions, with (1) weak connectivity and weak reciprocal connectivity, (2) strong connectivity but weak reciprocal connectivity, (3) weak connectivity but strong reciprocal connectivity, or strong connectivity and strong reciprocal connectivity. (C) Example dynamics of the mean synaptic variables *p* and *q*. Solid lines: theory, shaded lines: simulation. (D) Area of each dynamical region from panel A as the balance between potentiation and depression is varied.

### Homeostatic inhibitory STDP maintains stable activity during excitatory plasticity

In the previous section we focused on the evolution of excitatory synaptic weights through eSTDP. However, when the mean excitatory-excitatory weight *p* potentiates, positive feedback could in principle destabilize the network’s activity. Inhibitory synaptic plasticity was needed to homeostatically prevent runaway firing rate activity as *p* increased. The inhibitory STDP rule was parameterized by a homeostatic target excitatory rate 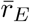 and if the excitatory rates were far from 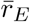 then the inhibitory STDP would become unbalanced and potentiation-dominated [Vogels et al., 2011]. In this case, the inhibitory plasticity would be dominated by firing rates so that:

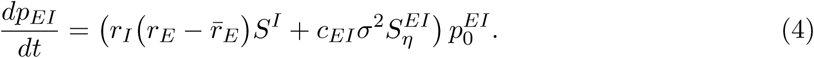

As before, *r*_*E*_ and *r*_*I*_ are the network-averaged excitatory and inhibitory firing rates. *c*_*EI*_ is the correlation of any external inputs targeting both excitatory and inhibitory cells, which we take to be 0. *S*^*I*^ is the area under the iSTDP rule and 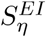 is the area under the iSTDP rule, weighted by the susceptibility of E-I neuron pairs to correlating inputs (Supporting Information S.1.1). If the excitatory rate was far from 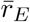, then *p*_*EI*_ and the firing rates *r*_*E*_, *r*_*I*_ evolved on a faster timescale than the balanced excitatory plasticity (Supporting Information S.1). The network thus had three timescales: the fast timescale of spiking correlations, the slow timescale of the homeostatic inhibitory STDP, and the slowest timescale of the balanced excitatory STDP.

The fixed points and stability of (*p*_*EI*_, *r*_*E*_, *r*_*I*_) on the unbalanced iSTDP timescale revealed that the inhibitory plasticity stabilizes the firing rates to keep 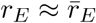 (more precisely to keep 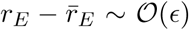; see Supporting Information S.1.3). Indeed, in simulations we saw that as *p* increased (decreased), *p*_*EI*_ potentiated (depressed) and maintained 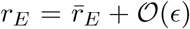 (Fig. 4A). The inhibitory-excitatory weight that maintained 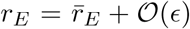 was the homeostatic fixed point, 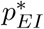.

The location of 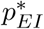 is given by solving the leading-order dynamics of the unbalanced inhibitory plasticity for the stable state: *dp*_*EI*_*/dt* = 0*, dr*_*E*_*/dt* = 0*, dr*_*I*_*/dt* = 0. This is a fixed point only on the timescale of the inhibitory STDP; as the excitatory-excitatory structure evolved, 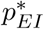 tracked the mean excitatory weight *p* to maintain stable activity (Fig. 4A). Due to the separation of timescales between the homeostatic iSTDP and the balanced eSTDP, we could predict the location of the homeostatic inhibitory weight 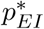 through a quasi-static approximation of *p* (Supporting Information S.1.3). We tracked the location of the homeostatic inhibitory weight 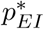 as a function of *p*. As expected, strong recurrent excitation required stronger inhibitory-excitatory feedback to enforce 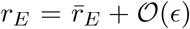 (Fig. 4B). In order to investigate the conditions under which inhibition was able to maintain stable activity at that homeostatic fixed point, we compared networks with plastic and non-plastic inhibition. With non-plastic inhibition, firing rates increased with *p*. If the excitatory feedback *p* became strong enough, the stationary firing rates lost stability (Fig. 4C). This instability was reflected in the development of hypersynchronous spiking, in contrast to the weakly correlated spiking activity in the network with plastic inhibition (Fig. 4D). In total, in order to study the robust potentiation of recurrent excitation, we required a counterbalancing potentiation of inhibitory onto excitatory neurons so as to homeostatically maintain a weakly correlated yet strongly connected excitatory network.

**Figure 4.**
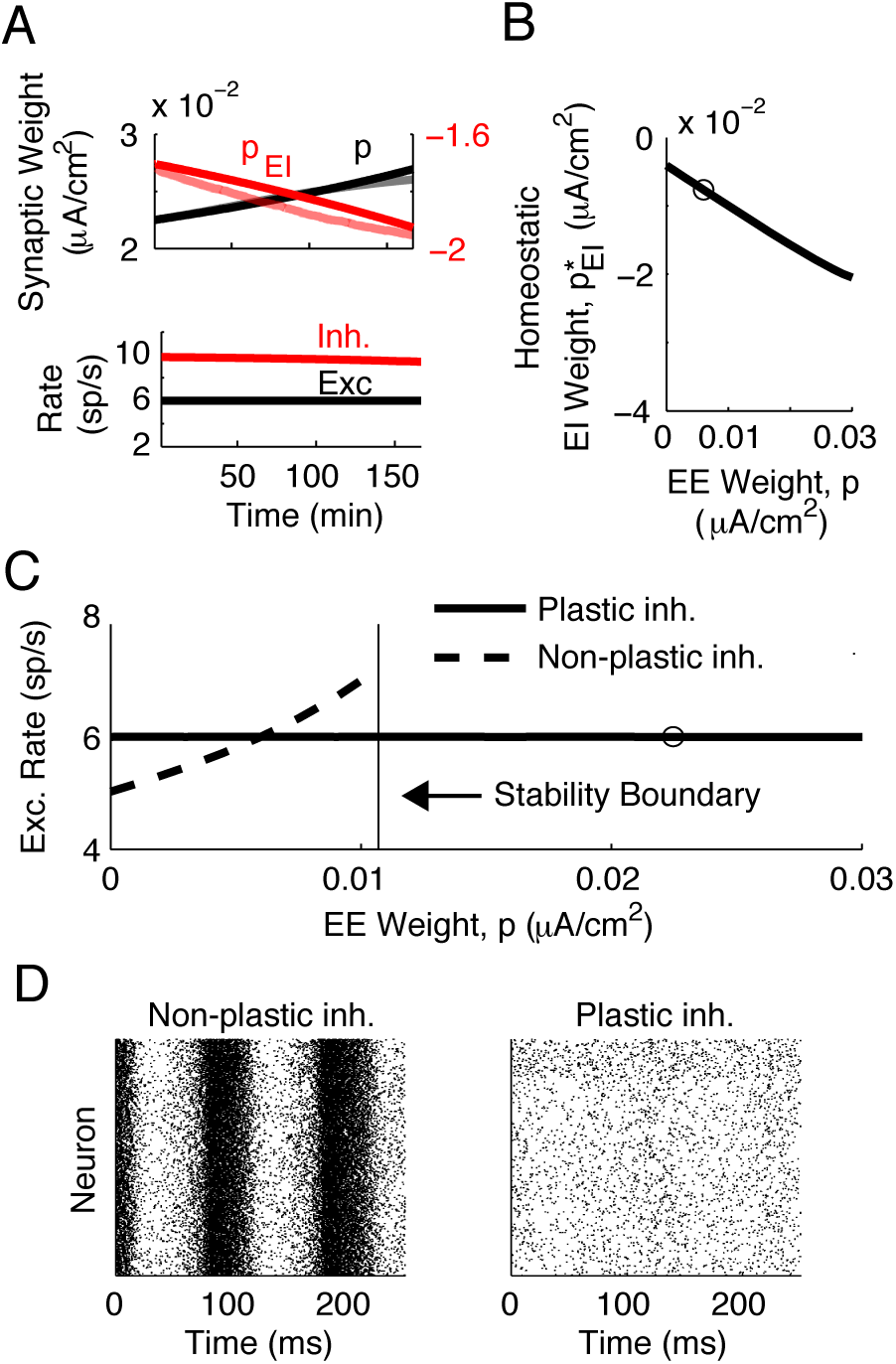
Inhibitory STDP maintains stable activity during excitatory plasticity. (A) Top: Coevolution of mean excitatory-excitatory synaptic weight *p* (black) and mean inhibitory-excitatory weight *p*_*EI*_ (red). Bottom: Firing rates during plasticity. (B) The homeostatic fixed point for *p*_*EI*_ as a function of the mean excitatory strength *p*. Open circle marks the inhibitory weight used for the nonplastic inhibition in C and D. (C) Firing rates as a function of excitatory weight in the cases of plastic and nonplastic inhibition. We predicted the location of that stability boundary by numerically computing the eigenvalues of the Fokker-Planck equation associated with the single-neuron voltage distribution and examining how activity is recurrently filtered through the network [Ledoux and Brunel, 2011]. (D) Raster plots of the network activity. In both bases the excitatory weight is at the value marked by the circle in panel D. For the right raster, *p*_*EI*_ is at its homeostatic fixed point.

### Stimulus-induced correlations drive assembly formation

The thresholds for potentiation and depression in both *p* and *q* suggested a mechanism for the formation of assembly structure through spike timing. Namely, if we define *p* and *q* variables for within- and cross-assembly connectivity, each should obey similar dynamics to Eqs. (2) and (3). In particular, each should have a threshold for potentiation. Furthermore, these thresholds should depend on the spatial correlation of the external inputs to within- or cross-cluster pairs of neurons. To test this, we divided the excitatory neurons into *M* putative assemblies of *κ* neurons each, based on their assigned stimulus preferences. Each assembly contained neurons that received spatially correlated inputs due to an external stimulus (Fig. 5A). The strength of these correlations was *c*_*AA*_ = 0.3; the correlation between inputs to neurons in different assemblies was *c*_*AB*_ = 0. In simulations, we saw that correlated external inputs drove the synaptic wiring into an assembly structure (Fig. 5B), which did not emerge spontaneously (Fig. 5C). We next sought a reduced mean field description of this plasticity. For ease of calculation, we assumed that the assemblies were symmetric so that the network structure was described by the four variables: (*p*_*AA*_, *p*_*AB*_, *q*_*AA*_, *q*_*AB*_) where *p*_*AA*_, *q*_*AA*_ describe within-assembly connectivity and *p*_*AB*_, *q*_*AB*_ describe cross-assembly connectivity:

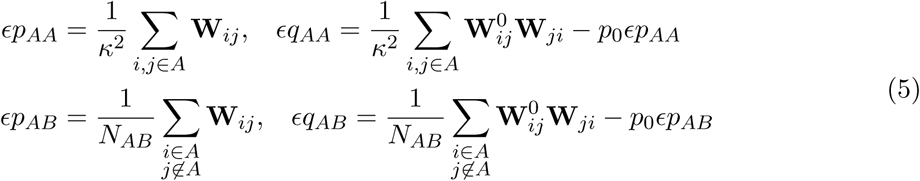

with *N*_*AB*_ = *κ* (*N*_*E*_ *− κ*) and now *ϵ* = 1*/*(*κp*_0_). (The definition of *ϵ* = 1*/*(*N*_*E*_*p*_0_) when discussing spontaneous plasticity above matches this definition if one considers the network of Figs. 2 & 3 to contain one excitatory cluster of *κ* = *N*_*E*_ neurons.) These measure the average strength of reciprocal connections either within (*q*_*AA*_) or between (*q*_*AB*_) assemblies, above what would be expected by chance. In our calculations we assumed that the assemblies were symmetric in that they had the same number of neurons and received inputs of the same strength, but did not assume that each neuron belonged to only one assembly. For simplicity, our simulations did have non-overlapping assemblies. Overlapping assemblies would mean that the correlation in external inputs to the different assemblies, *c*_*AB*_, should be greater than zero. Each of the mean field variables *p*_*AA*_, *p*_*AB*_, *q*_*AA*_, *q*_*AB*_ obeys dynamics of a similar form to those of *p, q* (see Supporting Information S.1) but the higher dimensional dynamics presents a complication for analysis.

**Figure 5.**
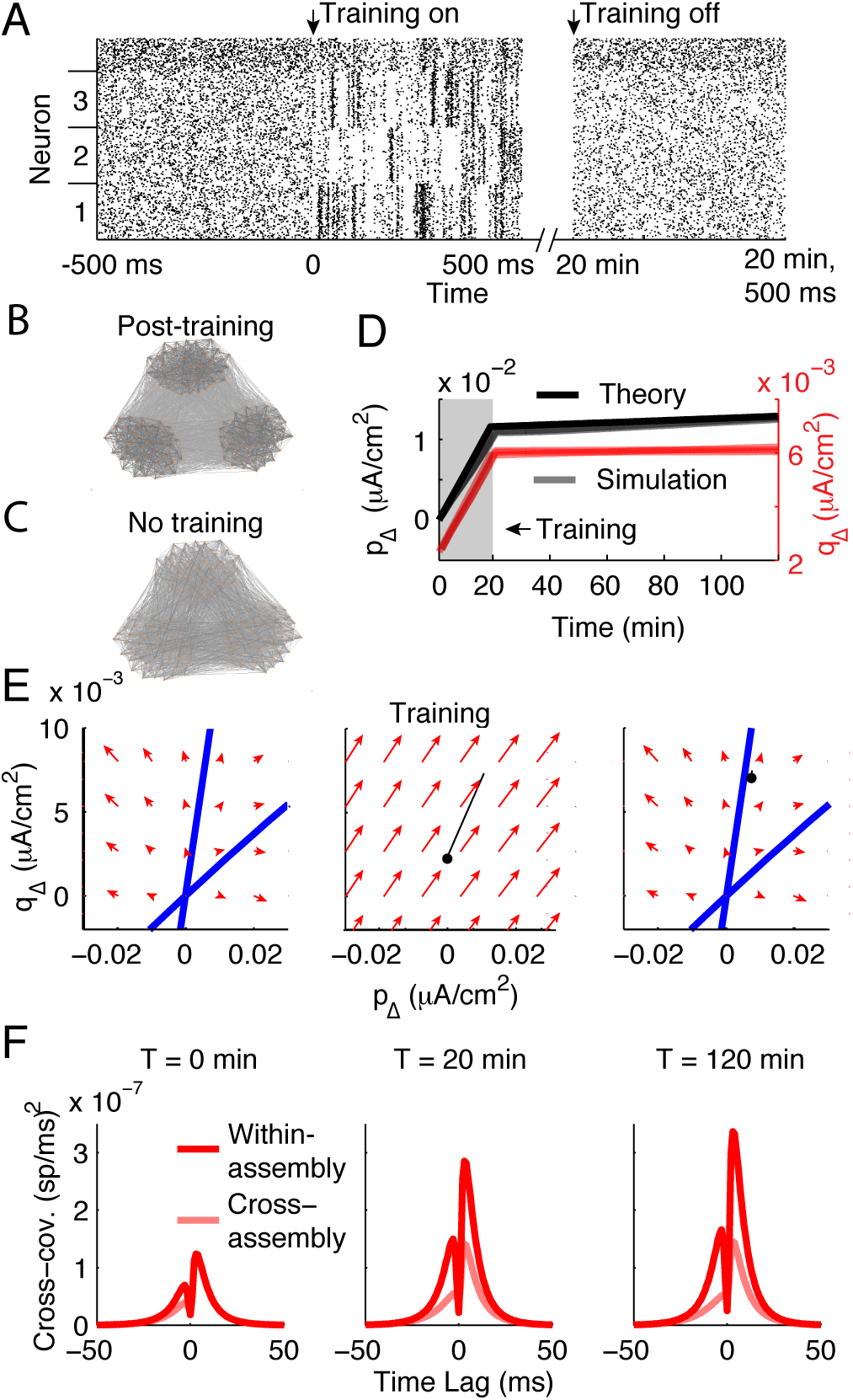
Common inputs promote self-reinforcing assemblies with strong reciprocal connectivity. (A) Raster of network activity during pre-training, training and post-training phases. Excitatory neurons ordered by assembly membership (labels on ordinate axis). (B,C) Visualization of connectivity between a subset of the excitatory neurons (50 from each of the three assemblies) after 180 minutes of simulated activity, with line darkness proportional to synaptic weight and nodes ordered by the Fruchterman-Reingold force algorithm. (D) Time course of the relative strength of within-assembly synapses, *p*_∆_ (black), and within-assembly reciprocal synapses, *q*_∆_ (red). (E) Phase plane of *p*_∆_*, q*_∆_ in the absence (pre- and post-training) or presence (training) of external input correlations. Blue: nullclines for *c*_∆_ = 0 were 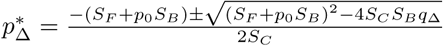 and 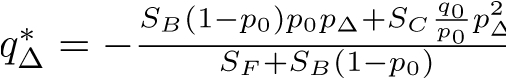. Black traces: simulation of the spiking network. (F) Average spike train cross-covariances (truncated approximation, Eq. S.10). Left, before training. Middle, immediately at end of training. Right, 100 min post-training.

In order to obtain a simpler description, we considered the change of variables:

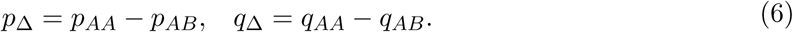

These measure the relative strength of assembly structure in the network, at the levels of mean connection strength (*p*_∆_) and above-chance reciprocal connection strength (*q*_∆_). For example, in order for a network to respect the structure observed in mouse V1 by Cossell et al. [Cossell et al., 2015], it should have *p*_∆_ > 0*, q*_∆_ > 0. The dynamics of (*p*_∆_, *q*_∆_) can be simply calculated from those of (*p*_*AA*_, *p*_*AB*_, *q*_*AA*_, *q*_*AB*_) and are (see Supporting Information S. 2):

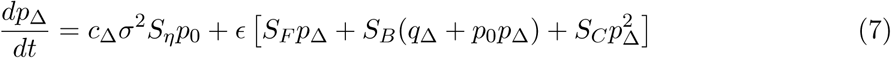

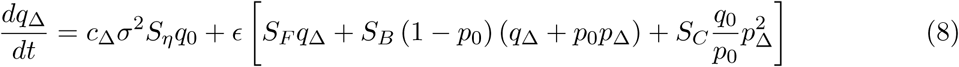

where *c*_∆_ = *c*_*AA*_ − *c*_*AB*_ is the relative strength of the external input correlation. Notably, the dynamics of *p*_∆_ and *q*_∆_ decoupled from the overall strengths of excitation and inhibition in the network; Eqs. (7) and (8) do not explictly depend on *p, q*. Further, the contribution of chance spike coincidences, 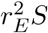, canceled because neurons in each assembly have the same average firing rate. Satisfyingly, the mean field theory of Eqs. (7) and (8) gave an accurate match to network simulations during training (*c*_∆_ > 0) and spontaneous (*c*_∆_ = 0) regimes (Figure 5D).

The dynamics of *p*_∆_ and *q*_∆_ have nullclines that represent thresholds for potentiation/depression (Fig. 5E, blue curves). The origin (*p*_∆_ = 0*, q*_∆_ = 0) is unstable and the nullclines divide the phase plane into four regions, containing each potential combination of potentiation and depression of (*p*_∆_*, q*_∆_). We take the synaptic weights to be initially unstructured, so that before training *p*_∆_ *≈ q*_∆_ *≈* 0 (Fig. 5E, left). If external input correlations are higher for within-assembly pairs than cross-assembly pairs (*c*_∆_ > 0), the unstable point at (0, 0) is shifted to negative (*p*_∆_*, q*_∆_) (Fig. 5E, middle). This pushed the unstable synaptic dynamics towards having assemblies of strongly reciprocally connected neurons. Furthermore, after the removal of the training signal the system had moved into a region of the phase plane where the assembly structure would reinforce itself (Fig. 5E, right). This occurred because the training of assembly structure gave rise to stronger spike-train correlations within assemblies than between (Fig. 5F left, middle). Indeed, the training of assembly structure into the network led to a doubling of spike train covariability for within-assembly neurons compared to cross-assembly neurons (Fig. 5F). This confirmed that the potentiation of (*p*_∆_*, q*_∆_) reflected potentiation of (*p*_*AA*_, *q*_*AA*_) rather than depression of *p*_*AB*_, *q*_*AB*_. In the absence of an over-representation of reciprocal connections, Ω = 0, *q*_*AA*_ and *q*_*AB*_ are negligible. In this case, the joint dynamics of *p*_*AA*_, *p*_*AB*_ are similar to those of *p*_∆_*, q*_∆_, also showing a threshold for assembly formation (Fig. S.2). If another set of assemblies were to be subsequently learned, then synapses would be reassigned from *p*_*AA*_, *q*_*AA*_ to *p*_*AB*_, *q*_*AB*_. Since the training dynamics promoted within-assembly connectivity rather than suppressing cross-assembly connectivity, we expect that a second set of assemblies could be learned without losing the first set of assemblies.

### Assembly training with the triplet STDP rule

So far, we have used a plasticity rule for excitatory synapses driven only by spike pairs. This type of rule fails to capture a key aspect of long-term plasticity: the plasticity rule’s dependence upon the overall firing rates of pre- and postsynaptic spike pairs (Fig. 1A). This simplification was necessary for the derivation of the mean-field theory in Eqs. (2), (3), (7), and (8). Nevertheless, it remains a concern that the results of the theory do not apply for more realistic plasticity rules. Thus motivated, we asked whether our results held in networks of model spiking neurons where the eSTDP rule depended upon spike time triplets [Pfister and Gerstner, 2006], so that firing rates could also shape the learning rule. Because we lacked a mean-field theory we restricted our analysis to numerical simulations of the network.

The triplet STDP rule has depression driven by spike pairs and potentiation driven by triplets of two post-synaptic spikes and one presynaptic spike (Fig. 6A; see Methods), with a balance between potentiation and depression so that uncorrelated pre- and postsynaptic spike trains would lead to no change in synaptic weight. Since spike triplet correlations were weaker than spike pair correlations, we assumed that within-assembly pairs of neurons received a baseline level of correlated external input (*c*_*AA*_ = 0.2). Without any additional training signal, the transient dynamics of the mean within- and cross-assembly weights exhibited three phases: 1) depression of both *p*_*AA*_ and *p*_*AB*_, 2) potentiation of *p*_*AA*_ once *p*_*AB*_ had decayed to 0, and 3) depression of *p*_*AA*_ to a weak baseline level (Fig. 6B, red). The mean inhibitory-excitatory synaptic weight *p*_*EI*_ exhibited similar dynamics, as did the mean reciprocal synaptic weights, *q*_*AA*_ and *q*_*AB*_ (Fig. 6C-D, red). The transitions between the first and second phases occurred when the cross-assembly connections had decayed to zero; the transition between the second and third reflected the more complicated dynamics of synaptic weights under triplet STDP. Even under this more complicated plasticity rule, the baseline input correlations were insufficient to drive strong assembly formation.

**Figure 6.**
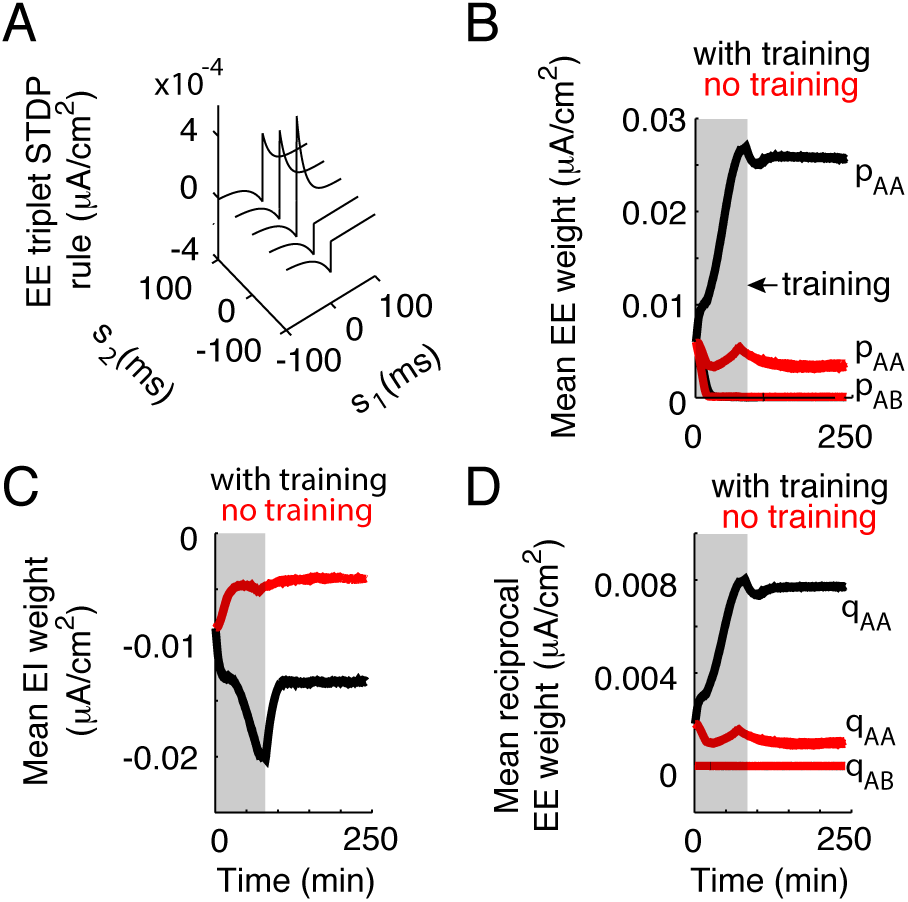
Spike time correlations can drive assembly formation and maintenance with triplet STDP. (A) Triplet STDP rule: the change in synaptic weight as a function of the pre-post time lag *s*_1_ and the time lag in between two postsynaptic spikes, *s*_2_. (B) Evolution of the mean synaptic weight within (*p*_*AA*_) and between (*p*_*AB*_) assemblies. (C) Evolution of the mean inhibitory-excitatory synaptic weight, *p*_*EI*_. (D) Evolution of the mean reciprocal synaptic weight within (*q*_*AA*_) and between (*q*_*AB*_) assemblies. (B-D) Black: with training (*c*_*AA*_ = 0.3 during the period highlighted in gray). Red: without training (*c*_*AA*_ = 0.2 always).

In order to drive assembly formation, we increased the input correlations to within-assembly pairs to the same strength as used with the pair-based STDP rule (*c*_*AA*_ = 0.3) for 133 minutes of simulation time (Fig. 6B-D, gray shaded area). This drove the formation of strong within-assembly connectivity and reciprocal connectivity (Fig. 6B-D, black) At the end of the training period, the external input correlations reverted to their baseline level of *c*_*AA*_ = 0.2. During the subsequent spontaneous activity, the trained assembly structure was stable: the mean within-assembly synaptic weight and reciprocal weight did not revert to their original levels (Fig. 6B-D, black). In contrast to the pair-based STDP rule, we did not see the trained structure spontaneously reinforce itself. The triplet STDP rule was additive (without weight-dependence), so we expect it to exhibit the unstable dynamics like the pair-based rule. However, triplet correlations are weaker than pairwise correlations when firing rates are low. A more complicated network structure could increase triplet correlations during spontaneous activity, making the dynamics of triplet STDP easier to observe [Jovanovi and Rotter, 2016].

### Trained noise covariance maintains coding performance

We finally asked how the spontaneous reinforcement of learned network structures, and the associated internally generated spike train covariability, affected the ability of cell assemblies to encode their preferred inputs. We took the partially symmetric network (Fig. 5) and allowed the external input to excitatory neurons in an assembly to depend on a stimulus *θ*: *µ*_E_ = *µ*_ext_ + *µ*_*θ*_. For simplicity, we took each stimulus to target exactly one assembly and considered only the coding by a single assembly (denoted *A*; Fig. 7A). We assumed that all neurons in assembly *A* had the same tuning. Although the stimulus only targeted one assembly, the value of *θ* varied continuously. In a visual analogy, *θ* could represent the contrast of a grating stimulus while its orientation would be reflected by the assembly it targeted. We measured the linear Fisher information, *FI*_*A*_(*θ*) [Beck et al., 2011] of an assembly’s total spike count 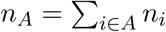 about the stimulus *θ*, where *n*_*i*_ is the spike count of neuron *i* (see Supporting Information S.4). Fisher information is a lower bound on the variance of any estimate of *θ* from *n*_*A*_, and the restriction to linear Fisher information gives a natural decomposition of *FI*_*A*_ into the response gain in *dn*_*A*_*/dθ* and response noise *c*_*AA*_. Since *n*_*A*_ naively sums the assembly activity we have that for large assemblies, the response variance 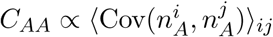, meaning that the mean pairwise covariance between neurons in an assembly is the dominant contribution to the noise in *n*_*A*_’s estimate of *θ*.

**Figure 7.**
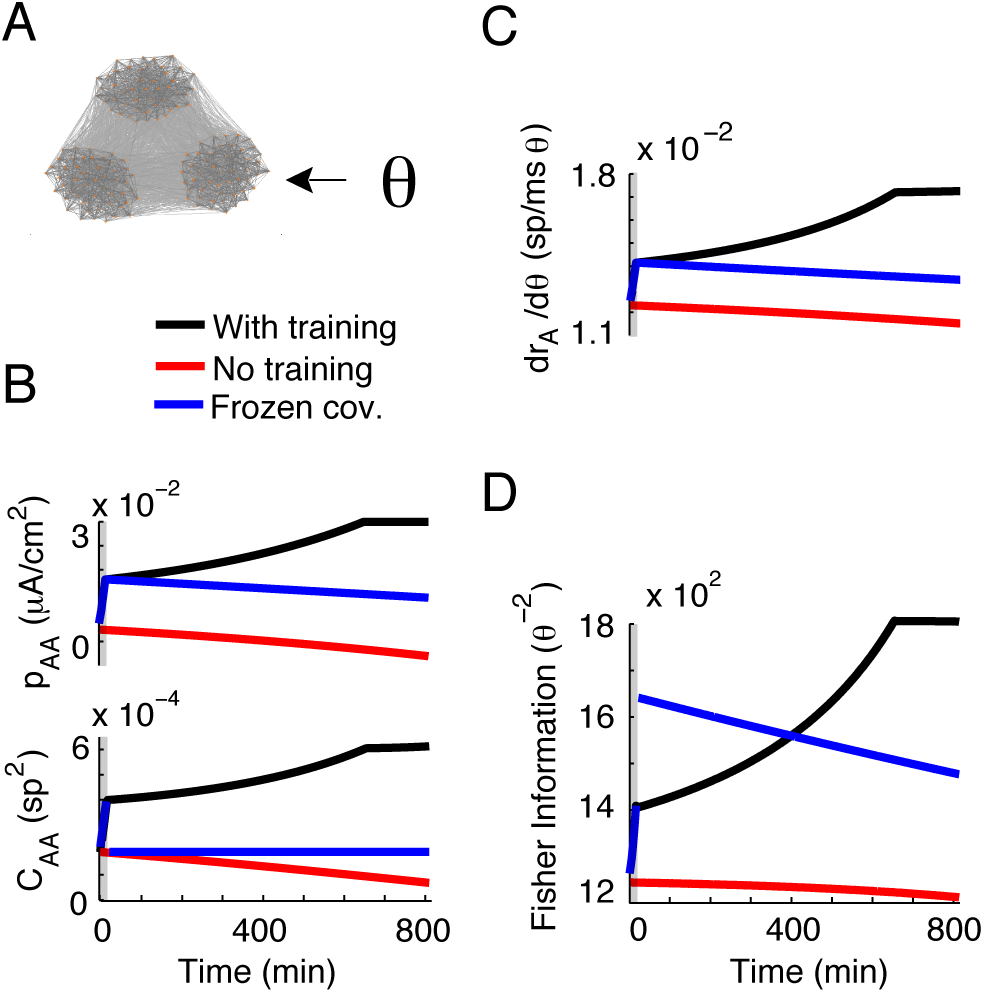
Internally generated spiking correlations reflect and reinforce assembly coding. (A) One assembly received a stimulus, *θ*, which it encodes by that assembly’s total spike count in *T* = 100 ms. (B) Top: Variance of the summed spike count of assembly *A* increased during and after training (black). Assembly *A*’s spike count variance decreased without training (red). As a control, we reset spiking covariability after training to pre-training values and froze them (blue). Bottom: The gain of stimulated neurons with respect to. *θ* increased during and after training (black). Without training, the stimulus-response gain decreased (red). With frozen covariances, the gain decreased after training (blue). Grey box: training period. (C) The mean strength of within-assembly connectivity. (D) Fisher information of the assembly’s spike count about *θ*.

We compared stimulus coding in networks with and without trained network structure (Fig. 7B top, black versus red curves). Since training increased spike train covariances for within-assembly pairs (Fig. 5F) networks, then the variance of the assembly’s summed spike count also increased with training and was reinforced after training (Fig. 7B bottom, black curve). In agreement, when the training signals were absent and assemblies did not form, then *C*_*AA*_ slowly decreased over time (Fig. 7B bottom, red curve).

One expectation from increased variability is that training assembly structure would, in our simplified coding scenario, be deleterious to stimulus coding since the internally generated correlations reflected the stimulus structure (with 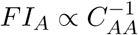) [Moreno-Bote et al., 2014]. To determine the net impact of training on *FI*_*A*_, we must evaluate how training and covariability affect the response gain which combines with *C*_*AA*_ to determine *FI*_*A*_. We calculated the stimulus-response gain *dn*_*A*_*/dθ*, taking into account direct stimulus-driven inputs and indirect filtering of the stimulus through recurrence onto assembly *A* (Supporting Information S.4). A consequence of trained assembly structure was increased gain through the positive feedback inherent within an assembly (Fig. 7C, black versus red curves). The increased gain outweighed the increased variability so that overall *FI*_*A*_ grew with training and further increased through the assembly reinforcement post-training (Fig. 7D, black versus red curves). In total, assembly formation was overall beneficial to network coding despite requiring larger overall network variability.

As a final illustration of the beneficial role of noise correlations for stimulus coding, we leveraged our theory in order to examine plasticity when spike train covariability was decoupled from connectivity. We considered an artificial scenario where training occurred and drove plasticity normally, yet immediately after training we reset the spike train covariability to its pre-training value and forced it to remain at this value (Fig. 7B, blue curves). We did this by freezing the right-hand side of Eqs. (7), (8) to its value given by the initial structure *p*_∆_(0)*, q*_∆_(0), with *c*_∆_ = 0. For a period of time post-training, the *FI*_*A*_ from this network with frozen covariances was larger than that of the trained network because of the combination of a large gain from training and low variability through the artificial reset (Fig. 7D, blue versus black curves). However, a consequence of low spike train covariability was a slow but clear degradation of assembly structure so that response gain reduced over time. This eventually reduced *FI*_*A*_ so that the network with internally generated covariability showed higher *FI*_*A*_ for times > 400 minutes after training. Thus, while noise correlations can have a detrimental impact on stimulus coding, the benefits of stimulus-specific recurrent structure and the role of spike train correlations play in maintaining that structure are such that noise correlations were beneficial in our simplified coding scenario.

## Discussion

### Rate-based versus timing–based assembly formation

Since early seminal work [Bell et al., 1997, Markram, 1997, Bi and Poo, 1998] there has been intense research on the role of spike timing in shaping synaptic strength [Markram et al., 2011, Markram et al., 2012]. Theoretical work with eSTDP in cortical networks first established the role of timing in the formation of feedforward structures [Masuda and Kori, 2007, Takahashi et al., 2009, Gerstner et al., 1996, Fiete et al., 2010, Tannenbaum and Burak, 2016]. More recently, eSTDP has been shown to promote the spontaneous formation of structured circuit motifs [Ocker et al., 2015] as well as support the stability of attractor network structure [Wei and Koulakov, 2014]. However, the role of spike timing in the formation of trained macroscopic assembly structure has been elusive. Past work in recurrent spiking network models has shown that the rate-dependence of eSTDP can be sufficient for forming and maintaining neuronal assembly structure [Litwin-Kumar and Doiron, 2014, Zenke et al., 2015]. Our study gives an alternative framework, where it is the fine-timescale correlations in spiking activity that drives assembly formation. We derived a low-dimensional mean field theory for the plasticity of neuronal assemblies in partially symmetric networks of integrate-and-fire neurons with excitatory STDP and homeostatic inhibitory STDP. This revealed that internally generated spike train correlations can provide a threshold for potentiation or depression of mean synaptic weight and for mean reciprocal connectivity. Spatial correlations in external inputs shifted these thresholds, promoting an assembly structure in the network. Furthermore, the post-training structure of spike train correlations reflected the learned network structure and actively reinforced that architecture. This promoted strong synaptic weights within assemblies and strong reciprocal connectivity within assemblies.

We used a particular strength of input correlation to entrain assembly structure, *c*_*AA*_ = 0.3, but our results are not particular to that parameter choice. Starting from a network without assembly structure (*p*_∆_ *≈ q*_∆_ *≈* 0), the training correlations need only to be sufficient to shift the thresholds (nullclines) for *p*_∆_ and *q*_∆_ below zero in order to push the network towards assembly structure. As long as during the training period, *p*_∆_ and *q*_∆_ pass the spontaneous locations of the nullclines given by *c*_∆_ = 0, then when the training signal is turned off the learned structure will self-reinforce (Fig. 5E). We cannot predict specific levels of feedforward input correlation to in vivo cortical assemblies because of this interdependence between the initial network structure and the strength of correlations necessary to entrain assembly structure.

The mechanisms behind rate-based and timing–based assembly formation are distinct. In the rate-based scenario, the training of assemblies is *sequential*—each stimulus is presented in the absence of other stimuli so that neuron pairs within the same assembly can have coordinated high firing rates to drive potentiation, while neuron pairs in different assemblies can have a high-low firing rates that drive depression. In the timing-based framework, assemblies can be trained in *parallel* since within- and cross-assembly neuron pairs can simultaneously receive correlated and uncorrelated external inputs. While both frameworks show a spontaneous reinforcement of assembly structure, the mechanics of reinforcement are quite different. In rate-based assembly formation the learned network structure is a stable *attractor* in the space of synaptic weights. Spontaneous reinforcement occurs if the network has not converged to the attractor during training [Litwin-Kumar and Doiron, 2014, Zenke et al., 2015]. If the network structure is perturbed from the attractor, then spontaneous activity will retrain the network [Litwin-Kumar and Doiron, 2014]. In our timing-based formation, it is the position of an unstable *repeller* that determines the growth or decay of structure. Spontaneous reinforcement occurs when the synaptic state is such that the repeller pushes dynamics towards assembly wiring.

While the mechanisms underlying rate- and timing-based assembly formation via eSTDP are distinct, they are not necessarily mutually exclusive. In mammals, visual stimuli lead to stimulus-specific changes in both firing rates and spike train correlations from the retina [Zylberberg et al., 2016] to the primary and higher visual cortices (e.g. [Kohn and Smith, 2005, Ito et al., 2010, Denman and Contreras, 2014, Ruff and Cohen, 2016, Ponce-Alvarez et al., 2013]), where they can shape the information carried about visual stimuli [Ohiorhenuan et al., 2010]. Spike time correlations and firing rate variations can both contribute to plasticity in more realistic plasticity models [Graupner et al., 2016], suggesting that the dynamics induced by these different forms of plasticity could be complementary, with spike time correlations driving plasticity to reinforce network structures from stimulus-induced firing rate-driven plasticity. Future work should investigate their interactions during learning and spontaneous activity, and in specified neural systems.

### Inhibitory plasticity and stabilization

Inhibitory feedback plays two main roles in this study. The first is to modulate excitatory plasticity by contributing to spike train covariability amongst excitatory neurons. The strength of this contribution is governed by the strength of the inhibitory feedback, which is in turn governed by inhibition’s second role: homeostatic control of firing rates.

In the absence of inhibition, potentiation of excitatory synapses in our networks led to runaway excitation, meaning that in the presence of inhibition the network existed in an inhibitory-stabilized regime (Figure **??**). In contrast to other recent studies [Litwin-Kumar and Doiron, 2014, Zenke et al., 2015], inhibitory STDP alone was sufficient to stabilize the network activity in our work without imposing synaptic scaling or other compensatory mechanisms. This was due to the relationship between the timescales of excitatory and inhibitory plasticity [Zenke and Gerstner, 2017]. We take excitatory plasticity to be balanced between potentiation and depression. This sets the dynamics of the mean excitatory weight *p* to occur on a slow timescale 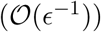 set by the eSTDP rule and the magnitude of spike train correlations. When the firing rates are maintained at their stable fixed points, the inhibitory STDP is similarly governed by a timescale set by the iSTDP rule and the magnitude of spike train correlations. If the firing rates are outside an 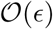 neigborhood around their fixed point, this causes the iSTDP rule to become unbalanced, so that it is governed by an 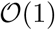 timescale (Supporting Information S.1). This feature—that the inhibitory STDP can become unbalanced in order to maintain stable activity—guarantees that it can dynamically stabilize the network activity in the face of the balanced excitatory plasticity. If, however, the amplitude of the iSTDP rule (the parameter *f*_*I*_) is sufficiently small 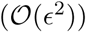, then the inhibitory STDP is recruited slowly and will not track the excitatory rates. In this case even if *r*_*E*_ were far from the homeostatic target rate 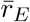, the leading-order dynamics of *p*_*EI*_ would be 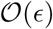; that is, the inhibitory STDP would be governed by the same timescale as the balanced eSTDP. Similarly, if the eSTDP did not have a balance between potentiation and depression, its leading-order dynamics would occur on an 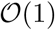 timescale with the homeostatic iSTDP - although these leading-order dynamics would not contain the 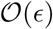 contributions of internally generated spike train correlations. In both of these cases, the joint dynamics of inhibitory and excitatory STDP could give rise to interesting interactions, and it remains to be shown whether these would be stable.

The stability of network activity and of learned synaptic strength are formally separate issues but are functionally very related to one another. Early studies of iSTDP in entorhinal cortex report a classic Hebbian asymmetric STDP curve [Haas et al., 2006], typically assumed for eSTDP (Fig. 2A). However, recent work in mouse auditory cortex report a symmetric iSTDP curve [Damour and Froemke, 2015], similar to that used in our study (Fig. 2B). Theoretical work has show that iSTDP with either asymmetric and symmetric learning curves will stabilize recurrent activity, provided that high postsynaptic activity recruits potentiation while low activity recruits depression [Luz and Shamir, 2012]. Thus, in our network iSTDP stabilizing recurrent excitation is likely not sensitive to the exact iSTDP curve used in our model. Nevertheless, homeostatic scaling of excitatory synapses, or other mechanisms for maintaining stable activity, would work in a similar fashion provided they occurred on a faster timescale than the balanced excitatory STDP. The relative timescales of Hebbian and homeostatic plasticity mechanisms and their joint dynamics are an active area of inquiry [Zenke and Gerstner, 2017].

The eSTDP rule we use models the induction of long-term plasticity, but does not account for longer-term maintenance or consolidation of synaptic resources. More complicated models explicitly include these mechanisms, but are less amenable to mathematical analysis [Clopath et al., 2008, Higgins et al., 2014]. Alternatively, simple STDP models with weight-dependent synaptic updates (multiplicative) models or in which the potentiation window is shifted so that pre-post pairs at small lags cause depression can yield unimodal weight distributions [Rubin et al., 2001, Van Rossum et al., 2000, Babadi and Abbott, 2010]. When these models have unbalanced potentiation and depression so that the plasticity is dominated by chance spike coincidences, we expect these to exhibit stable equilibria for the mean-field synaptic weight variables [Lajoie et al., 2017]. The dynamics of intrinsically stable STDP models with balanced potentiation and depression, on the other hand, are less obvious.

The question of how neurons can undergo associative, Hebbian learning while maintaining stable activity has long been studied [Miller and MacKay, 1994, Miller, 1996]. While homeostasis is often thought of as a slower process than learning, recent work has highlighted the necessity of homeostatic mechanisms operating on a comparable timescale to excitatory plasticity [Zenke et al., 2013]. Homeostatic regulation acting alone, however, can paradoxically destabilize network activity, inducing oscillations in neurons’ firing rates [Harnack et al., 2015]. Homeostatic regulation mediated by diffusive transmitters like nitrous oxide can have different effects than that mediated by synaptic mechanisms [Sweeney et al., 2015].

The study of how homeostatic regulation and mechanisms for associative learning interact to allow stable memories and stable activity remains an exciting area of open inquiry [Keck et al., 2017].

### Correlated spontaneous activity can maintain coding performance

Many theoretical studies have asked how the joint trial-to-trial fluctuations in population response (noise correlations) impact population coding [Averbeck et al., 2006, Kohn et al., 2016]. The mechanisms behind noise correlations are varied [Doiron et al., 2016], and both feedforward [Kanitscheider et al., 2015] and recurrent [Helias et al., 2014] circuits can contribute to linking stimulus and noise correlations. When noise and signal correlations have similar enough structures, noise correlations generally impair coding [Moreno-Bote et al., 2014, Hu et al., 2014, Zylberberg et al., 2016, Franke et al., 2016]. In the absence of sensory stimulation, patterns of activity across cortical populations are often similar to those observed during sensory stimulation [Arieli et al., 1996, Luczak et al., 2009, Xu et al., 2012]. Thus, the circuits that support correlated variability in spontaneous states likely overlap with the circuits responsible for noise correlations in evoked state.

Neuron pairs with similar tuning often show larger noise correlations than pairs with dissimilar tuning [Aertsen et al., 1989, Ahissar et al., 1992, Espinosa and Gerstein, 1988, Kohn and Smith, 2005, Ko et al., 2011, Cossell et al., 2015], suggesting that the circuit dynamics giving rise to them would impair coding. Spontaneous activity is also usually viewed as a problem for plasticity: learned weight changes must be stable in the face of spontaneous activity. We demonstrated a novel and complementary viewpoint on the impact of noise correlations on population coding which belies these hypotheses.

In our study, the trained network architecture produced sizable within-assembly spontaneous correlations that combined with the STDP rule to reinforce assembly structure. If the assembly wiring was originally due to a shared stimulus input, then the spontaneous correlations needed to retain structure will be a source of noise correlations when the stimulus is to be coded. Our simplified stimulus coding scenario was such that within-assembly noise correlations degraded the neural code. However, the strong positive feedback from within-assembly recurrence enhanced the response gain, which improved coding. We suggest that noise correlations could be a mechanism for the active maintenance of assembly structure and the high response gain it confers to a neuronal population. This finding does not critically depend on the fast-timescale coordinated spiking activity required for STDP, and stability of assembly structure through long-timescale firing rate correlations should have a similar effect [Litwin-Kumar and Doiron, 2014, Wei and Koulakov, 2014]. How this finding interacts with distributed population codes, and how plasticity distributes information amongst different dimensions of population activity, remains to be investigated.

## Materials and Methods

A detailed description of the derivation of the mean field plasticity dynamics for *p, q* and *p*_*AA*_, *p*_*AB*_, *q*_*AA*_, *q*_*AB*_ and *p*_∆_*, q*_∆_ is given in Supporting Information S.1 and S.2. A detailed description of the derivation of the plasticity of the Fisher information is given in Supporting Information S.4.

### 1 Network model

We consider a network of *N* neurons, *N*_*E*_ of which are excitatory and divided into *M* clusters of size *κ*. There are *N*_*I*_ = *γκ* inhibitory neurons. Model parameters are in Table 1. In order to reflect the overrepresentation of reciprocally connected pairs of excitatory neurons in cortex [Song et al., 2005, Perin et al., 2011], we took the baseline excitatory-excitatory connectivity of our network, 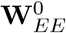, to be composed of two parts: a symmetric random binary matrix, 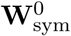 with connection probability Ω*p*_0_, and an asymmetric random binary matrix 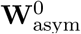 with connection probability (1 − Ω)*p*_0_. Both had Erős-Rènyi statistics. (Note that here, an asymmetric matrix does not mean that there are no reciprocal synapses, but rather that connections *i → j* and *i ← j* are uncorrelated; reciprocal connections occur in 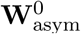 with frequency ((1 − Ω)*p*_0_)^2^.) The parameter Ω determined the frequency of bidirectionally connected pairs of excitatory neurons in 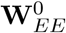, while *p*_0_ sets the overall connection probability. We took Ω = 0.4, which with *p*_0_ = .15 yielded networks with bidirectional connections approximately 3 times as frequent as they would be with Ω = 0, comparable to that reported in rat sensory cortices [Song et al., 2005, Perin et al., 2011]. Excitatory-inhibitory, inhibitory-excitatory and inhibitory-inhibitory connectivity were asymmetric, with a connection probability of 0.4, reflecting the relatively dense inhibitory connectivity observed in cortex [Hofer et al., 2011, Fino and Yuste, 2011]. We remark that the direct evidence for Ω > 0 is restricted to the visual and somatosensory cortices of young mice; it remains to be shown more broadly across species and into adulthood. Additionally we exclude autapses 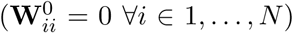. This means that the excitatory-excitatory connectivity is characterized by its empirical connection density *p*_0_ and the frequency of loops *q*_0_

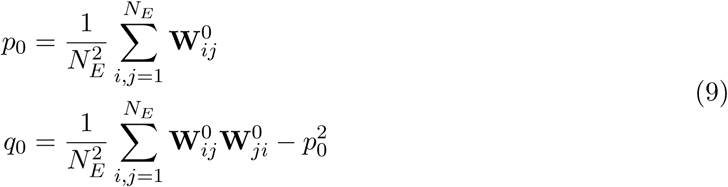

**Table 1.** Neuron and synapse parameters

| Parameter | Description | Value |
| --- | --- | --- |
| $C$ | Membrane capacitance | $1 \mu\text{F}/\text{cm}^2$ |
| $g_L$ | Leak conductance | $0.1\text{mS}/\text{cm}^2$ |
| $V_L$ | Leak reversal potential | $-72 \text{ mV}$ |
| $\Delta$ | Action potential steepness | $1.4 \text{ mV}$ |
| $V_T$ | Action potential initiation threshold | $-48 \text{ mV}$ |
| $V_{th}$ | Action potential threshold | $30 \text{ mV}$ |
| $V_{re}$ | Action potential reset | $-72 \text{ mV}$ |
| $\tau_{ref}$ | Action potential width | $2 \text{ ms}$ |
| $\mu$ | External input mean | $1 \mu\text{A}/\text{cm}^2$ |
| $\sigma$ | External input standard deviation | $9 \text{ mV}$ |
| $W^{\text{max},\text{E}}$ | Maximum synaptic weight | $15\epsilon \mu\text{A}/\text{cm}^2$ |
| $W^{\text{max},\text{I}}$ | Maximum synaptic weight | $-7.5\epsilon \mu\text{A}/\text{cm}^2$ |
| $J(t)$ | Synaptic filter (EPSC shape) | $\exp(-(t/\tau_s))$ |
| $\tau_{\text{sE}}$ | Excitatory synaptic time constant | $2 \text{ ms}$ |
| $\tau_{\text{sI}}$ | Inhibitory time constant | $10 \text{ ms}$ |

We assume that the statistics of the adjacency matrix for within- and between-assembly connectivity are the same (and equal to *p*_0_ and *q*_0_). The synaptic weight matrix, **W**, is initially generated from **W**^0^ by giving each synapse the same initial weight. We consider the mean strength of E-E synapses within one cluster *A* and from other clusters into cluster *A*, *p*_*AA*_ and *p*_*AB*_ respectively:

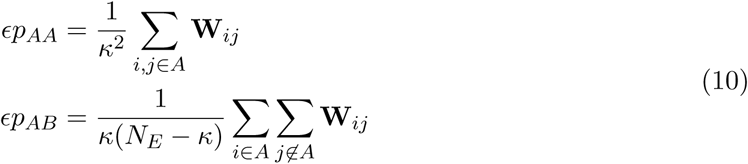

The small parameter ε= (*κp*_0_)^−1^ scales the synaptic weights. We take the mean strength of connections within each cluster to be symmetric and the strength of connections into any one cluster from outside to be the same as into the others (so for all clusters *A* and *B*, *p*_*AA*_ = *p*_*BB*_ and *p*_*AB*_ = *p*_*BA*_). Similarly, we measure the strength of *reciprocal* connections within a cluster, *q*_*AA*_, or between clusters, *q*_*AB*_:

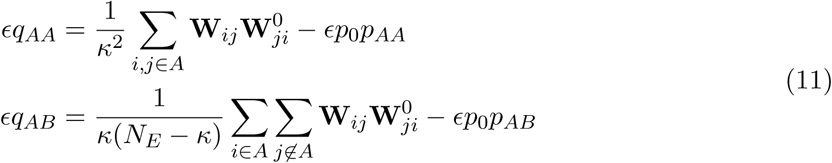

By subtracting off *p*_0_*p*_*AA*_ in the definition of *q*_*AA*_ (and likewise for *q*_*AB*_), we measure the mean strength of reciprocal connections above what would be expected in a network with no correlations between synapses. Note: if the network is asymmetric (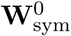 = **0**) then *q*_0_ is negligible 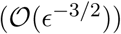 and so are the initial values of *q*_*AA*_ and *q*_*AB*_.

We take the connectivity in between inhibitory and excitatory neurons, and within inhibitory neurons, to have (asymmetric) Erdős-Rényi statistics, so that these are characterized by their mean synaptic weights: *p*_*EI*_ for inhibitory *→* excitatory connections,

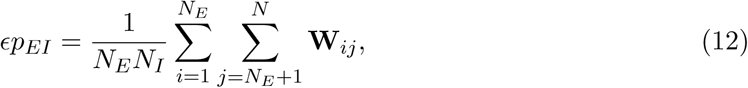

and likewise *p*_*IE*_ and *p*_*II*_.

Finally, individual neurons had exponential integrate-and-fire (EIF) dynamics [Fourcaud-Trocme et al., 2003], part of a class of models well-known to capture the spike initiation dynamics of cortical neurons [Jolivet et al., 2004, Jolivet et al., 2008]. Neurons’ membrane voltages obeyed:

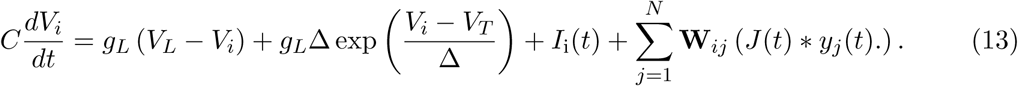

The external input decomposes as *I*_*i*_(*t*) = *µ* + *σξ*_*i*_(*t*), where *ξ*_*i*_(*t*) is a white noise term described by 〈*ξ*_*i*_(*t*)〉 = 0 and 〈*ξ*_*i*_(*t*)*ξ*_*j*_(*t′*)〉 = *δ_ij_δ*(*t − t′*) with *δ*_*ij*_ being a Kronecker delta function while *δ*(*t*) is a Dirac delta function. In order to impose correlated external inputs, we decomposed *ξ*_*i*_(*t*) as 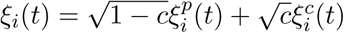 where 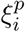 and 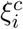 are the private and common parts of the external input to that neuron, respectively. *c* depended on the joint assembly membership of each pair of neurons so that for *M* assemblies, there were *M* different common external training inputs. For *i* ≠ *j*, we have 〈*ξ*_*i*_(*t*)*ξ*_*j*_(*t′*)〉 = *cδ*(*t − t′*). The network model parameters are given in Table 1.

## 2 Plasticity models

### 2.1 Excitatory Plasticity

Synapses between excitatory neurons undergo additive Hebbian eSTDP:

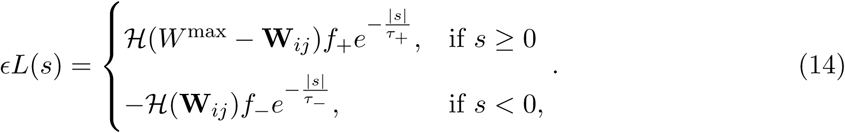

where *s* = *t*_*post*_ − *t*_*pre*_ is the time lag between spikes. *f*_*±*_ give the amplitude of individual changes in synaptic weights due to potentiation (*f*_+_) or depression (*f*_*−*_), and the time constants *τ*_*±*_ determine how synchronous spike pairs must be to cause plasticity. When an excitatory postsynaptic neuron spikes at time *t*_*i*_, the weights of the excitatory synapses onto it are updated as 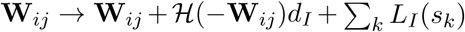, where *s*_*k*_ = *t*_*i*_ − *t*_*k*_ is the time difference between that postsynaptic spike a previous presynaptic spike (at time *t*_*k*_) of neuron *j*. The parameters of the eSTDP model are in Table 2.

**Table 2.** Plasticity model parameters

| Parameter | Description | Value |
| --- | --- | --- |
| $\tau_-$ | Time constant of eSTDP depression window | $30 \text{ ms}$ |
| $\tau_+$ | Time constant of eSTDP potentiation window | $15 \text{ ms}$ |
| $S$ | Integral of eSTDP rule | $-W^{\text{max},\text{E}}/4000 \mu\text{A}/\text{cm}^2 \text{ ms}$ |
| $f_-$ | Amplitude of eSTDP depression window | $-W^{\text{max},\text{E}}/1000 \mu\text{A}/\text{cm}^2 \text{ ms}$ |
| $f_+$ | Amplitude of eSTDP potentiation window | Given by $\delta$ and $f_-$, $\tau_+$, $\tau_-$ |
| $\tau_I$ | Time constant of iSTDP rule | $15\text{ms}$ |
| $f_I$ | Amplitude of iSTDP rule | $W^{\text{max},\text{I}}/1000$ |
| $\bar{r}_E$ | Homeostatic target E rate | $6 \text{ sp/ms}$ |

We also tested a triplet-based eSTDP model (Fig. 6). In this model, excitatory-excitatory synapses potentiated due to triplets of two postsynaptic spikes and one presynaptic spike:

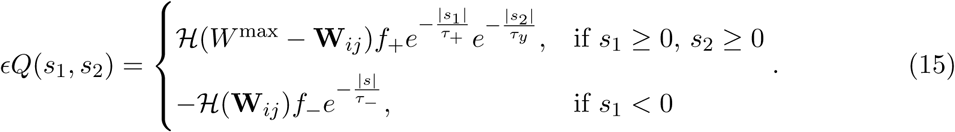

where *s*_1_ is the pre-post time lag and *s*_2_ the post-post time lag. This yields potentiation due to pre-post-post spike triplets and post-pre-post spike triplets and depression due to post-pre spike pairs. This model was introduced as a minimal model for fitting V1 slice data by [Pfister and Gerstner, 2006]. Parameters of the triplet eSTDP model are in Table 3.

**Table 3.** Triplet STDP parameters

| Parameter | Description | Value |
| --- | --- | --- |
| $\tau_-$ | Time constant of eSTDP depression window | 33.7 ms |
| $\tau_+$ | Time constant for pre-post potentiation | 16.8 ms |
| $\tau_y$ | Time constant for post-post potentiation | 114 ms |
| $S$ | Integral of triplet STDP rule | $0 \mu\text{A}/\text{cm}^2 \text{ ms}$ |
| $f_-$ | Amplitude of depression window | $-W^{\text{max,E}}/1000 \mu\text{A}/\text{cm}^2 \text{ ms}$ |
| $f_+$ | Amplitude of potentiation window | Given by $S$ and $f_-$, $\tau_+$, $\tau_-$, $\tau_y$ |

### 2.2 Inhibitory Plasticity

In recurrent networks, excitatory plasticity can lead to the destabilization of asynchronous activity [Lubenov and Siapas, 2008] and the development of pathological synchrony [Morrison et al., 2007]. Past modeling studies have explored plasticity of inhibition as a stabilizing mechanism, preventing runaway activity in networks with [Litwin-Kumar and Doiron, 2014, Zenke et al., 2015] and without [Vogels et al., 2011] excitatory plasticity. Following past studies [Vogels et al., 2011, Litwin-Kumar and Doiron, 2014, Zenke et al., 2015] the inhibitory *→* excitatory synapses obey the iSTDP rule:

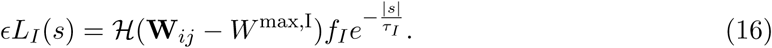

In addition to this pair-based rule, each presynaptic (inhibitory) spike drives depression of the inhibitory synapses by 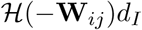 where 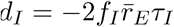. When an excitatory neuron spikes at time *t*_*i*_, the weights of the inhibitory synapses onto it are updated as 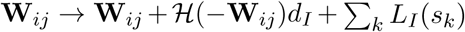, where *s*_*j*_ = *t*_*i*_ − *t*_*k*_ is the time difference between that postsynaptic spike and a presynaptic spike at time *t*_*k*_.

The iSTDP rule acts as a form of homeostatic control where pairs of near coincident pre- and postsynaptic spikes caused potentiation of inhibitory-excitatory synapses, while individual presynaptic spikes caused depression [Vogels et al., 2011] (Figure **??**A). The strength of this depression was determined by the homeostatic target excitatory rate, 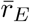. The iSTDP model parameters are given in Table 2.

## Acknowledgments

We thank Ken Miller, Bard Ermentrout, Jonathan Rubin, Anne-Marie Oswald, Thanos Tzounopoulos and Krešimir Josić for their comments on an early draft of this manuscript, and the anonymous referees for the comments which greatly improved the manuscript.

## References

Aertsen et al., 1989. Aertsen, A., Gerstein, G., Habib, M., and Palm, G. (1989). Dynamics of neuronal firing correlation: modulation of ”effective connectivity”. J Neurophysiol, 61:900–917.

Ahissar et al., 1992. Ahissar, E., Vaadia, E., Ahissar, M., Bergman, H., Arieli, A., and Abeles, M. (1992). Dependence of cortical plasticity on correlated activity of single neurons and on behavioral context. Science (New York, N.Y.), 257(5075):1412–1415.

Arieli et al., 1996. Arieli, A., Sterkin, A., Grinvald, A., and Aertsen, A. (1996). Dynamics of ongoing activity: explanation of the large variability in evoked cortical responses. Science (New York, N.Y.), 273(5283):1868–1871.

Averbeck et al., 2006. Averbeck, B. B., Latham, P. E., and Pouget, A. (2006). Neural correlations, population coding and computation. Nat Rev Neurosci, 7(5):358–66.

Babadi and Abbott, 2010. Babadi, B. and Abbott, L. F. (2010). Intrinsic Stability of Temporally Shifted Spike-Timing Dependent Plasticity. PLoS Comput Biol, 6(11):e1000961.

Bair et al., 2001. Bair, W., Zohary, E., and Newsome, W. T. (2001). Correlated firing in macaque visual area MT: time scales and relationship to behavior. J Neurosci, 21:1676–1697.

Beck et al., 2011. Beck, J., Bejjanki, V. R., and Pouget, A. (2011). Insights from a simple expression for linear fisher information in a recurrently connected population of spiking neurons. Neural Computation, 23(6):1484–1502.

Bell et al., 1997. Bell, C. C., Han, V. Z., Sugawara, Y., and Grant, K. (1997). Synaptic plasticity in a cerebellum-like structure depends on temporal order. Nature, 387(6630):278–281.

Bi and Poo, 1998. Bi, G.-q. and Poo, M.-m. (1998). Synaptic modifications in cultured hippocampal neurons: dependence on spike timing, synaptic strength, and postsynaptic cell type. The Journal of Neuroscience, 18(24):10464–10472.

Buzski, 2010. Buzski, G. (2010). Neural syntax: cell assemblies, synapsembles, and readers. Neuron, 68(3):362–385.

Brgers et al., 2012. Brgers, C., Franzesi, G. T., LeBeau, F. E. N., Boyden, E. S., and Kopell, N. J. (2012). Minimal Size of Cell Assemblies Coordinated by Gamma Oscillations. PLOS Computational Biology, 8(2):e1002362.

Carrillo-Reid et al., 2016. Carrillo-Reid, L., Yang, W., Bando, Y., Peterka, D. S., and Yuste, R. (2016). Imprinting and recalling cortical ensembles. Science, 353(6300):691–694.

Clopath et al., 2008. Clopath, C., Ziegler, L., Vasilaki, E., Bsing, L., and Gerstner, W. (2008). Tag-Trigger-Consolidation: A Model of Early and Late Long-Term-Potentiation and Depression. PLoS Comput Biol, 4(12):e1000248.

Clopath, Claudia et al., 2010. Clopath, Claudia, Bsing, Lars, Vasilaki, Eleni, and Gerstner, Wulfram (2010). Connectivity reflects coding: a model of voltage-based STDP with homeostasis. Nat Neurosci, 13(3):344–352.

Cossell et al., 2015. Cossell, L., Iacaruso, M. F., Muir, D. R., Houlton, R., Sader, E. N., Ko, H., Hofer, S. B., and Mrsic-Flogel, T. D. (2015). Functional organization of excitatory synaptic strength in primary visual cortex. Nature, 518(7539):399–403.

Denman and Contreras, 2014. Denman, D. J. and Contreras, D. (2014). The structure of pairwise correlation in mouse primary visual cortex reveals functional organization in the absence of an orientation map. Cerebral Cortex (New York, N.Y.: 1991), 24(10):2707–2720.

Doiron et al., 2016. Doiron, B., Litwin-Kumar, A., Rosenbaum, R., Ocker, G. K., and Josi, K. (2016). The mechanics of state-dependent neural correlations. Nature Neuroscience, 19(3):383–393.

Damour and Froemke, 2015. Damour, J. and Froemke, R. (2015). Inhibitory and Excitatory Spike-Timing-Dependent Plasticity in the Auditory Cortex. Neuron, 86(2):514–528.

Espinosa and Gerstein, 1988. Espinosa, I. E. and Gerstein, G. L. (1988). Cortical auditory neuron interactions during presentation of 3-tone sequences: effective connectivity. Brain Research, 450(1-2):39–50.

Feldman, 2012. Feldman, D. (2012). The Spike-Timing Dependence of Plasticity. Neuron, 75(4):556–571.

Fiete et al., 2010. Fiete, I. R., Senn, W., Wang, C. Z. H., and Hahnloser, R. H. R. (2010). Spike-Time-Dependent Plasticity and Heterosynaptic Competition Organize Networks to Produce Long Scale-Free Sequences of Neural Activity. Neuron, 65(4):563–576.

Fino and Yuste, 2011. Fino, E. and Yuste, R. (2011). Dense inhibitory connectivity in neocortex. Neuron, 69(6):1188–203.

Fourcaud-Trocme et al., 2003. Fourcaud-Trocme, N., Hansel, D., van Vreeswijk, C., and Brunel, N. (2003). How spike generation mechanisms determine the neuronal response to fluctuating inputs. Journal of Neuroscience, 23(37):11628–11640.

Franke et al., 2016. Franke, F., Fiscella, M., Sevelev, M., Roska, B., Hierlemann, A., and AzeredodaSilveira, R. (2016). Structures of Neural Correlation and How They Favor Coding. Neuron, 89(2):409–422.

Gerstner et al., 1996. Gerstner, W., Kempter, R., van Hemmen, J. L., and Wagner, H. (1996). A neuronal learning rule for sub-millisecond temporal coding. Nature, 383(6595):76–78.

Gerstner et al., 1993. Gerstner, W., Ritz, R., and van Hemmen, J. L. (1993). Why spikes? Hebbian learning and retrieval of time-resolved excitation patterns. Biological Cybernetics, 69(5-6):503–515.

Graupner and Brunel, 2012. Graupner, M. and Brunel, N. (2012). Calcium-based plasticity model explains sensitivity of synaptic changes to spike pattern, rate, and dendritic location. Proceedings of the National Academy of Sciences, 109(10):3991–3996.

Graupner et al., 2016. Graupner, M., Wallisch, P., and Ostojic, S. (2016). Natural Firing Patterns Imply Low Sensitivity of Synaptic Plasticity to Spike Timing Compared with Firing Rate. Journal of Neuroscience, 36(44):11238–11258.

Haas et al., 2006. Haas, J. S., Nowotny, T., and Abarbanel, H. D. I. (2006). Spike-Timing-Dependent Plasticity of Inhibitory Synapses in the Entorhinal Cortex. Journal of Neurophysiology, 96(6):3305–3313.

Harnack et al., 2015. Harnack, D., Pelko, M., Chaillet, A., Chitour, Y., and van Rossum, M. C. (2015). Stability of Neuronal Networks with Homeostatic Regulation. PLoS Comput Biol, 11(7):e1004357.

Harris and Mrsic-Flogel, 2013. Harris, K. D. and Mrsic-Flogel, T. D. (2013). Cortical connectivity and sensory coding. Nature, 503(7474):51–58.

Hebb, 1949. Hebb, D. O. (1949). The organization of behavior: a neuropsychological theory. L. Erlbaum Associates, Mahwah, N.J.

Helias et al., 2014. Helias, M., Tetzlaff, T., and Diesmann, M. (2014). The Correlation Structure of Local Neuronal Networks Intrinsically Results from Recurrent Dynamics. PLoS Comput Biol, 10(1):e1003428.

Higgins et al., 2014. Higgins, D., Graupner, M., and Brunel, N. (2014). Memory Maintenance in Synapses with Calcium-Based Plasticity in the Presence of Background Activity. PLoS Comput Biol, 10(10):e1003834.

Hofer et al., 2011. Hofer, S. B., Ko, H., Pichler, B., Vogelstein, J., Ros, H., Zeng, H., Lein, E., Lesica, N. A., and Mrsic-Flogel, T. D. (2011). Differential connectivity and response dynamics of excitatory and inhibitory neurons in visual cortex. Nature Neuroscience, 14(8):1045–1052.

Hu et al., 2014. Hu, Y., Zylberberg, J., and Shea-Brown, E. (2014). The Sign Rule and Beyond: Boundary Effects, Flexibility, and Noise Correlations in Neural Population Codes. PLoS Comput Biol, 10(2):e1003469.

Ito et al., 2010. Ito, H., Maldonado, P., and Gray, C. (2010). Dynamics of Stimulus-Evoked Spike Timing Correlations in the Cat Lateral Geniculate Nucleus. Neuroscience Research, 68:e74.

Jia et al., 2013. Jia, X., Tanabe, S., and Kohn, A. (2013). Gamma and the Coordination of Spiking Activity in Early Visual Cortex. Neuron, 77(4):762–774.

Jolivet et al., 2004. Jolivet, R., Lewis, T. J., and Gerstner, W. (2004). Generalized Integrate-and-Fire Models of Neuronal Activity Approximate Spike Trains of a Detailed Model to a High Degree of Accuracy. Journal of Neurophysiology, 92(2):959–976. We demonstrate that single-variable integrate-and-fire models can quantitatively capture the dynamics of a physiologically detailed model for fast-spiking cortical neurons. Through a systematic set of approximations, we reduce the conductance-based model to 2 variants of integrate-and-fire models. In the first variant (nonlinear integrate-and-fire model), parameters depend on the instantaneous membrane potential, whereas in the second variant, they depend on the time elapsed since the last spike [Spike Response Model (SRM)]. The direct reduction links features of the simple models to biophysical features of the full conductance-based model. To quantitatively test the predictive power of the SRM and of the nonlinear integrate-and-fire model, we compare spike trains in the simple models to those in the full conductance-based model when the models are subjected to identical randomly fluctuating input. For random current input, the simple models reproduce 7080 percent of the spikes in the full model (with temporal precision of 2 ms) over a wide range of firing frequencies. For random conductance injection, up to 73 percent of spikes are coincident. We also present a technique for numerically optimizing parameters in the SRM and the nonlinear integrate-and-fire model based on spike trains in the full conductance-based model. This technique can be used to tune simple models to reproduce spike trains of real neurons.

Jolivet et al., 2008. Jolivet, R., Schrmann, F., Berger, T. K., Naud, R., Gerstner, W., and Roth, A. (2008). The quantitative single-neuron modeling competition. Biological Cybernetics, 99(4-5):417–426.

Jovanovi and Rotter, 2016. Jovanovi, S. and Rotter, S. (2016). Interplay between Graph Topology and Correlations of Third Order in Spiking Neuronal Networks. PLOS Comput Biol, 12(6):e1004963.

Kanitscheider et al., 2015. Kanitscheider, I., Coen-Cagli, R., and Pouget, A. (2015). Origin of information-limiting noise correlations. Proceedings of the National Academy of Sciences, 112(50):e6973–e6982.

Keck et al., 2017. Keck, T., Toyoizumi, T., Chen, L., Doiron, B., Feldman, D. E., Fox, K., Gerstner, W., Haydon, P. G., Hbener, M., Lee, H.-K., Lisman, J. E., Rose, T., Sengpiel, F., Stellwagen, D., Stryker, M. P., Turrigiano, G. G., and Rossum, M. C. v. (2017). Integrating Hebbian and homeostatic plasticity: the current state of the field and future research directions. Phil. Trans. R. Soc. B, 372(1715):20160158.

Kempter et al., 1999. Kempter, R., Gerstner, W., and Van Hemmen, J. L. (1999). Hebbian learning and spiking neurons. Physical Review E, 59(4):4498.

Kim et al., 2016. Kim, T., Oh, W. C., Choi, J. H., and Kwon, H.-B. (2016). Emergence of functional subnetworks in layer 2/3 cortex induced by sequential spikes in vivo. Proceedings of the National Academy of Sciences, 113(10):e1372–e1381.

Ko et al., 2013. Ko, H., Cossell, L., Baragli, C., Antolik, J., Clopath, C., Hofer, S. B., and Mrsic-Flogel, T. D. (2013). The emergence of functional microcircuits in visual cortex. Nature, 496(7443):96–100.

Ko et al., 2011. Ko, H., Hofer, S. B., Pichler, B., Buchanan, K. A., Sjstrm, P. J., and Mrsic-Flogel, T. D. (2011). Functional specificity of local synaptic connections in neocortical networks. Nature, 473(7345):87–91.

Ko et al., 2014. Ko, H., Mrsic-Flogel, T. D., and Hofer, S. B. (2014). Emergence of Feature-Specific Connectivity in Cortical Microcircuits in the Absence of Visual Experience. The Journal of Neuroscience, 34(29):9812–9816.

Kohn et al., 2016. Kohn, A., Coen-Cagli, R., Kanitscheider, I., and Pouget, A. (2016). Correlations and Neuronal Population Information. Annual Review of Neuroscience, 39(1):237–256.

Kohn and Smith, 2005. Kohn, A. and Smith, M. A. (2005). Stimulus Dependence of Neuronal Correlation in Primary Visual Cortex of the Macaque. The Journal of Neuroscience, 25(14):3661–3673.

Lajoie et al., 2017. Lajoie, G., Krouchev, N. I., Kalaska, J. F., Fairhall, A. L., and Fetz, E. E. (2017). Correlation-based model of artificially induced plasticity in motor cortex by a bidirectional brain-computer interface. PLOS Computational Biology, 13(2):e1005343.

Ledoux and Brunel, 2011. Ledoux, E. and Brunel, N. (2011). Dynamics of networks of excitatory and inhibitory neurons in response to time-dependent inputs. Frontiers in Computational Neuroscience, 5:25.

Lee et al., 2016. Lee, W.-C. A., Bonin, V., Reed, M., Graham, B. J., Hood, G., Glattfelder, K., and Reid, R. C. (2016). Anatomy and function of an excitatory network in the visual cortex. Nature, 532(7599):370–374.

Litwin-Kumar and Doiron, 2014. Litwin-Kumar, A. and Doiron, B. (2014). Formation and maintenance of neuronal assemblies through synaptic plasticity. Nature Communications, 5.

Lubenov and Siapas, 2008. Lubenov, E. V. and Siapas, A. G. (2008). Decoupling through synchrony in neuronal circuits with propagation delays. Neuron, 58(1):118–131.

Luczak et al., 2009. Luczak, A., Barth, P., and Harris, K. D. (2009). Spontaneous Events Outline the Realm of Possible Sensory Responses in Neocortical Populations. Neuron, 62(3):413–425.

Luz and Shamir, 2012. Luz, Y. and Shamir, M. (2012). Balancing Feed-Forward Excitation and Inhibition via Hebbian Inhibitory Synaptic Plasticity. PLoS Comput Biol, 8(1):e1002334.

Markram, 1997. Markram, H. (1997). Regulation of Synaptic Efficacy by Coincidence of Postsynaptic APs and EPSPs. Science, 275(5297):213–215.

Markram et al., 2011. Markram, H., Gerstner, W., and Sjstrm, P. J. (2011). A history of spike-timing-dependent plasticity. Frontiers in Synaptic Neuroscience, 3:4.

Markram et al., 2012. Markram, H., Gerstner, W., and Sjstrm, P. J. (2012). Spike-timing-dependent plasticity: a comprehensive overview. Frontiers in Synaptic Neuroscience, page 2.

Masuda and Kori, 2007. Masuda, N. and Kori, H. (2007). Formation of feedforward networks and frequency synchrony by spike-timing-dependent plasticity. Journal of Computational Neuroscience, 22(3):327–345.

Miller, 1996. Miller, K. D. (1996). Synaptic economics: competition and cooperation in synaptic plasticity. Neuron, 17(3):371–374.

Miller and MacKay, 1994. Miller, K. D. and MacKay, D. J. C. (1994). The Role of Constraints in Hebbian Learning. Neural Computation, 6(1):100–126.

Mongillo et al., 2005. Mongillo, G., Curti, E., Romani, S., and Amit, D. J. (2005). Learning in realistic networks of spiking neurons and spike-driven plastic synapses. The European Journal of Neuroscience, 21(11):3143–3160.

Moreno-Bote et al., 2014. Moreno-Bote, R., Beck, J., Kanitscheider, I., Pitkow, X., Latham, P., and Pouget, A. (2014). Information-limiting correlations. Nature Neuroscience, 17(10):1410–1417.

Morrison et al., 2007. Morrison, A., Aertsen, A., and Diesmann, M. (2007). Spike-Timing-Dependent Plasticity in Balanced Random Networks. Neural Computation, 19(6):1437–1467.

Neves et al., 2008. Neves, G., Cooke, S. F., and Bliss, T. V. P. (2008). Synaptic plasticity, memory and the hippocampus: a neural network approach to causality. Nature Reviews Neuroscience, 9(1):65–75.

Ocker et al., 2015. Ocker, G. K., Litwin-Kumar, A., and Doiron, B. (2015). Self-Organization of Microcircuits in Networks of Spiking Neurons with Plastic Synapses. PLoS Comput Biol, 11(8):e1004458.

Ohiorhenuan et al., 2010. Ohiorhenuan, I. E., Mechler, F., Purpura, K. P., Schmid, A. M., Hu, Q., and Victor, J. D. (2010). Sparse coding and high-order correlations in fine-scale cortical networks. Nature, 466(7306):617–621.

Perin et al., 2011. Perin, R., Berger, T. K., and Markram, H. (2011). A synaptic organizing principle for cortical neuronal groups. Proceedings of the National Academy of Sciences, 108(13):5419–5424.

Pfister and Gerstner, 2006. Pfister, J.-P. and Gerstner, W. (2006). Triplets of Spikes in a Model of Spike Timing-Dependent Plasticity. The Journal of Neuroscience, 26(38):9673–9682.

Ponce-Alvarez et al., 2013. Ponce-Alvarez, A., Thiele, A., Albright, T. D., Stoner, G. R., and Deco, G. (2013). Stimulus-dependent variability and noise correlations in cortical MT neurons. Proceedings of the National Academy of Sciences, 110(32):13162–13167.

Rothschild et al., 2010. Rothschild, G., Nelken, I., and Mizrahi, A. (2010). Functional organization and population dynamics in the mouse primary auditory cortex. Nature Neuroscience, 13(3):353–360.

Rubin et al., 2001. Rubin, J., Lee, D., and Sompolinsky, H. (2001). Equilibrium Properties of Temporally Asymmetric Hebbian Plasticity. Physical Review Letters, 86(2):364–367.

Ruff and Cohen, 2016. Ruff, D. A. and Cohen, M. R. (2016). Stimulus Dependence of Correlated Variability across Cortical Areas. Journal of Neuroscience, 36(28):7546–7556.

Salkoff et al., 2015. Salkoff, D. B., Zagha, E., Yzge,., and McCormick, D. A. (2015). Synaptic Mechanisms of Tight Spike Synchrony at Gamma Frequency in Cerebral Cortex. Journal of Neuroscience, 35(28):10236–10251.

Sjstrm et al., 2001. Sjstrm, P. J., Turrigiano, G. G., and Nelson, S. B. (2001). Rate, timing, and cooperativity jointly determine cortical synaptic plasticity. Neuron, 32(6):1149–1164.

Song et al., 2000. Song, S., Miller, K. D., and Abbott, L. F. (2000). Competitive Hebbian learning through spike-timing-dependent synaptic plasticity. Nature Neuroscience, 3(9):919–926.

Song et al., 2005. Song, S., Sjstrm, P. J., Reigl, M., Nelson, S., and Chklovskii, D. B. (2005). Highly nonrandom features of synaptic connectivity in local cortical circuits. PLoS Biol, 3(3):e68.

Sweeney et al., 2015. Sweeney, Y., Hellgren Kotaleski, J., and Hennig, M. H. (2015). A Diffusive Homeostatic Signal Maintains Neural Heterogeneity and Responsiveness in Cortical Networks. PLoS Comput Biol, 11(7):e1004389.

Takahashi et al., 2009. Takahashi, Y. K., Kori, H., and Masuda, N. (2009). Self-organization of feed-forward structure and entrainment in excitatory neural networks with spike-timing-dependent plasticity. Physical Review E, 79(5):051904.

Tannenbaum and Burak, 2016. Tannenbaum, N. R. and Burak, Y. (2016). Shaping Neural Circuits by High Order Synaptic Interactions. PLOS Comput Biol, 12(8):e1005056.

van Rossum et al., 2000. van Rossum, M. C., Bi, G. Q., and Turrigiano, G. G. (2000). Stable Hebbian learning from spike timing-dependent plasticity. The Journal of Neuroscience, 20(23):8812–8821.

Vogels et al., 2011. Vogels, T. P., Sprekeler, H., Zenke, F., Clopath, C., and Gerstner, W. (2011). Inhibitory Plasticity Balances Excitation and Inhibition in Sensory Pathways and Memory Networks. Science, 334(6062):1569–1573.

Wei and Koulakov, 2014. Wei, Y. and Koulakov, A. A. (2014). Long-Term Memory Stabilized by Noise-Induced Rehearsal. Journal of Neuroscience, 34(47):15804–15815.

Xu et al., 2012. Xu, S., Jiang, W., Poo, M.-m., and Dan, Y. (2012). Activity recall in a visual cortical ensemble. Nature Neuroscience, 15(3):449–455.

Yoshimura et al., 2005. Yoshimura, Y., Dantzker, J. L. M., and Callaway, E. M. (2005). Excitatory cortical neurons form fine-scale functional networks. Nature, 433(7028):868–873.

Zenke et al., 2015. Zenke, F., Agnes, E. J., and Gerstner, W. (2015). Diverse synaptic plasticity mechanisms orchestrated to form and retrieve memories in spiking neural networks. Nature Communications, 6.

Zenke and Gerstner, 2017. Zenke, F. and Gerstner, W. (2017). Hebbian plasticity requires compensatory processes on multiple timescales. Phil. Trans. R. Soc. B, 372(1715):20160259.

Zenke et al., 2013. Zenke, F., Hennequin, G., and Gerstner, W. (2013). Synaptic Plasticity in Neural Networks Needs Homeostasis with a Fast Rate Detector. PLoS Comput Biol, 9(11):e1003330.

Zylberberg et al., 2016. Zylberberg, J., Cafaro, J., Turner, M. H., Shea-Brown, E., and Rieke, F. (2016). Direction-Selective Circuits Shape Noise to Ensure a Precise Population Code. Neuron, 89(2):369–383.

